# Task complexity temporally extends the causal requirement for visual cortex in perception

**DOI:** 10.1101/2021.06.22.449366

**Authors:** Matthijs N. Oude Lohuis, Jean L. Pie, Pietro Marchesi, Jorrit S. Montijn, Christiaan P.J. de Kock, Cyriel M. A. Pennartz, Umberto Olcese

## Abstract

The transformation of sensory inputs into behavioral outputs is characterized by an interplay between feedforward and feedback operations in cortical hierarchies. Even in simple sensorimotor transformations, recurrent processing is often expressed in primary cortices in a late phase of the cortical response to sensory stimuli. This late phase is engaged by attention and stimulus complexity, and also encodes sensory-independent factors, including movement and report-related variables. However, despite its pervasiveness, the nature and function of late activity in perceptual decision-making remain unclear. We tested whether the function of late activity depends on the complexity of a sensory change-detection task. Complexity was based on increasing processing requirements for the same sensory stimuli. We found that the temporal window in which V1 is necessary for perceptual decision-making was extended when we increased task complexity, independently of the presented visual stimulus. This window overlapped with the emergence of report-related activity and decreased noise correlations in V1. The onset of these co-occurring activity patterns was time-locked to and preceded reaction time, and predicted the reduction in behavioral performance obtained by optogenetically silencing late V1 activity (>200 ms after stimulus onset), a result confirmed by a second multisensory task with different requirements. Thus, although early visual response components encode all sensory information necessary to solve the task, V1 is not simply relaying information to higher-order areas transforming it into behavioral responses. Rather, task complexity determines the temporal extension of a loop of recurrent activity, which overlaps with report-related activity and determines how perceptual decisions are built.

## Introduction

During perceptual decision making, stimulus presentation triggers an early response component in primary sensory cortices (driven by thalamic bottom-up input^1^) and, often, a late component, thought to mostly result from recurrent activity through top-down, cross-areal interactions^2–4^. Traditional accounts of how sensory stimuli are transformed into appropriate behavioral outputs have mostly characterized this process in terms of feedforward architectures, where progressively higher-order areas extract sensory features of increasing complexity^5^ to eventually instruct motor output. In the visual cortical system, a fast-acting (<150 ms) feedforward sweep is sufficient for image categorization^6^. Accordingly, deep feedforward neural networks, inspired by this cortical hierarchical architecture, achieve near-human performance in image recognition^7,8^. The function of recurrent architectures has been primarily interpreted in the context of processing ambiguous or complex stimuli, for cognitive processes such as attention, and for consciousness^9–12^. For example, extra-classical receptive field effects in the visual system, such as surround suppression, and separating objects from background, are thought to depend on feedback projections from higher to lower visual areas^13–16^. Perceptual decisions involving figure-ground segregation require recurrent processing^13^, the duration of which becomes longer as a function of visual scene complexity^17^. Recently, a form of late activity in rodent V1 that reflects non-sensory variables such as movement, perceptual report, and arousal^18–23^ was shown to originate in prefrontal areas and progressively involve more posterior areas including sensory cortices^18,22^.

Many hypotheses have been proposed on the function of late, recurrent activity in sensory cortices (including distributed motor command generation and context-dependent sensory processing)^19^, but how it causally contributes to perception is debated. Across primates and rodents, the magnitude of late activity correlates with behavioral reports of perception^3,4,24,25^. Suppressing late activity in the primary somatosensory cortex impairs tactile detection^25^, whereas in primary visual cortex it has been argued that feedforward activity is sufficient for visual discrimination^6,26^. We hypothesize that the relative complexity of a task, which is captured by the set of task rules as instantiated in an attentional set – see e.g.^27^ – determines whether late activity plays a causal role in perceptual decision making, independently of stimulus complexity. For instance, integration with other sensory modalities^12,28,29^ may also extend the time required by frontal and pre-motor regions to converge to a decision. This process might reflect an evidence accumulation model^30,31^, i.e. a need to integrate information originating in V1 for longer periods in the case of complex, multisensory tasks. Analogously, the predictive processing framework^15,32,33^ posits that visual and decision-related areas will keep on interacting via recurrent connections to jointly represent sensory stimuli and transform them into appropriate motor responses, performing computations for a time interval that may depend on task complexity. Therefore, task complexity may extend the temporal window during which V1 activity remains causally relevant for perception, independently of visual stimulus features.

## Results

To address this hypothesis, we trained mice in three versions of an audiovisual change detection task (task A) with the same stimulus configurations, but different reward contingencies. Head-fixed mice were presented with a continuous audiovisual stream of inputs (Fig. 1a), with occasional instantaneous changes in the orientation of the drifting grating (visual trial) or the center frequency and harmonics of a Shepard Tone^34^ (auditory trial, Fig. 1b,c). We varied the amount of orientation (visual saliency) and frequency change (auditory saliency) across each animal’s perceptual threshold and fit all behavioral data according to a psychometric multi-alternative signal detection framework^35^.

**Figure 1:**
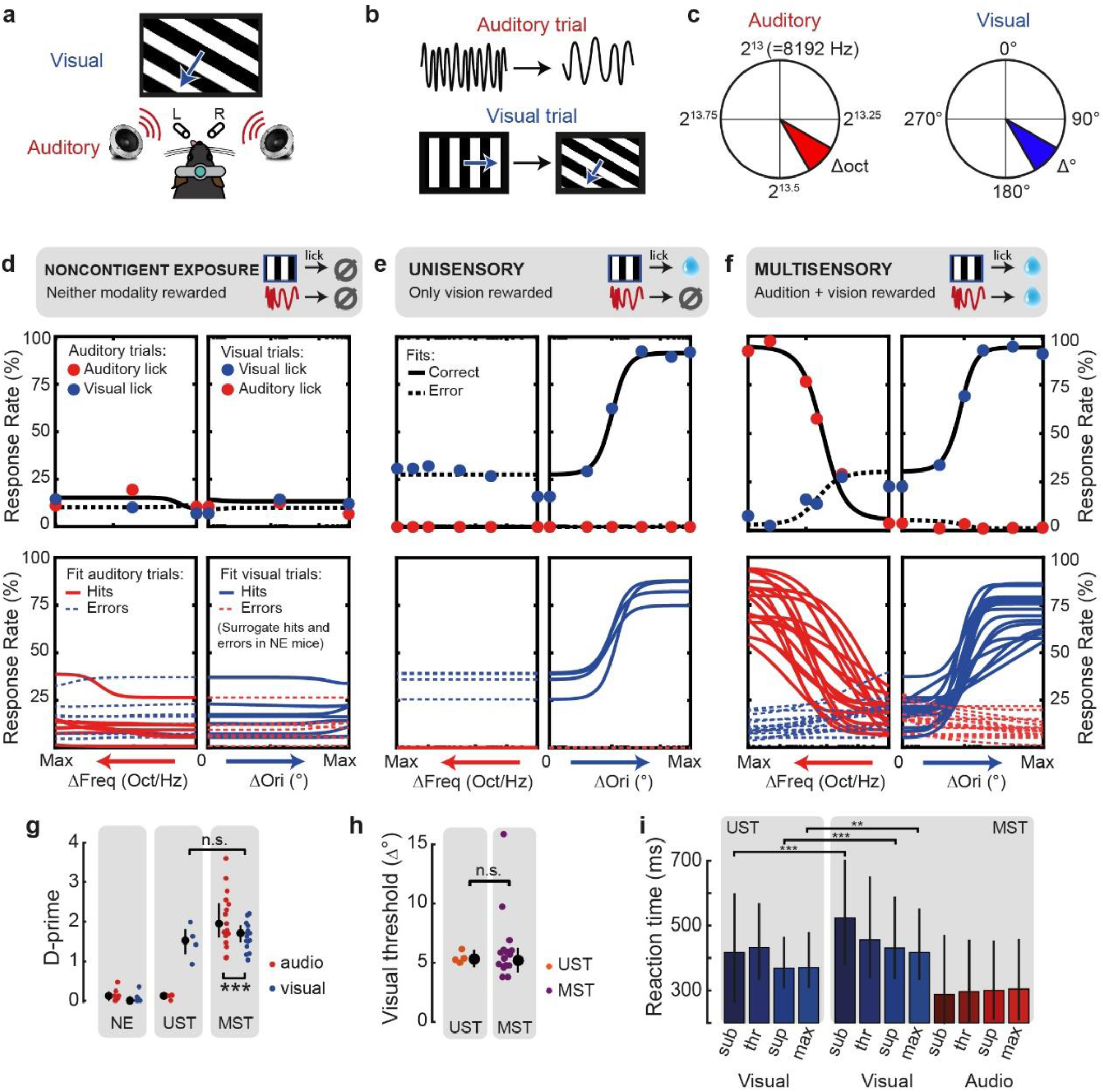
Multisensory task contingencies delay reaction time. a) In task A, head-fixed mice were presented with auditory and visual stimuli and two spouts (L: left and R: right) that detected licks and delivered rewards. b) Stimuli were continuous drifting gratings and Shepard tones. In auditory trials, the compound frequency of the Shepard tone changed, while in visual trials the orientation changed. c) Auditory center tone was defined in scientific pitch and changed in partial octaves (2^13^ Hz corresponds to C_9_, 2^13.25^ corresponds to D_9#_). The amount of frequency (left, red) or orientation (right, blue) change (small vs. large) determined saliency. Two units of auditory change were used (see Ext. Data Fig. 1). d) We trained three cohorts of mice on different reward contingencies with these same stimuli. Noncontingently exposed (NE) mice were rewarded for licks during response windows that were temporally decorrelated from the sensory stimuli. Post-hoc, licks to the visual spout and auditory spout that happened to fall after a stimulus change were defined as surrogate ‘hits’ and ‘errors’ to compare conditions across cohorts (see Methods). The upper panels show behavioral response rates (dots) and model fits (lines) for an example session. The bottom panels show the average psychometric fits for each mouse obtained by averaging parameters over sessions. Each session was fit with a two-alternative signal detection model (black lines in upper panels, colored in lower panels). e) Same as (d), but for UST animals, i.e. trained on vision only. Note how visual hit rates increase as a function of the amount of visual change, but not auditory change. The relatively high error rate to auditory changes arises because of licks to the visual spout, which was the only side associated with reward in this task. f) Same as (d), but for MST animals trained on both audition and vision (lick left for auditory, lick right for visual). Hit rates increased as a function of both visual and auditory change. g) The d-prime at maximum saliency for auditory (red) and visual (blue) trials across cohorts. Visual d-prime was comparable for UST and MST (p=0.3). Auditory d-prime was significantly higher than visual d-prime (p=1.86e-6). Each dot is the average over sessions for each animal. h) The detection threshold for visual orientation changes was comparable for UST and MST (p=0.87). Each dot is the average over sessions for one animal. i) Reaction time for the same subjectively salient visual stimuli (see Methods) was significantly shorter for UST compared to MST for 3 out of 4 saliency levels. Saliency levels: sub=subthreshold, thr=threshold, sup=suprathreshold, max=maximal change. Error bars denote the median and interquartile ranges. **p<0.01, ***p<0.001

We implemented three distinct task contingencies. First, for noncontingently exposed mice (*NE*, n=7 mice) neither vision nor audition was predictive of reward, and these mice did not selectively respond to the stimuli (Fig. 1d). In a second version, only vision was associated with reward, and these unisensory-trained mice (*UST*, n=4) were thus trained to selectively respond to visual changes only, and ignore auditory changes (Fig. 1e). Third, multisensory-trained mice (*MST*, n=17) were trained to detect both visual and auditory changes (Fig. 1f, e.g. lick left for vision, lick right for audition). Phrased differently, all mice were presented with the same stimuli during training and testing, but lick responses to visual changes were only rewarded in UST and MST mice, and auditory changes only in MST mice. To compare across cohorts, we also defined (surrogate) ‘hit’ and ‘miss’ trials for NE mice, based on whether (unrewarded) licks were performed after stimulus change. In all cohorts, mice performed many trials (mean 569, range 210-1047 per session). The discriminability index (d-prime) was high only for rewarded contingencies, in both the auditory and visual modality (Fig. 1g, for individual mice, see Ext. Data Fig. 1).

### Multisensory task contingencies delay reaction time

First, we wondered if visual performance was similar in the unisensory and multisensory task variants (UST and MST) and whether the more complex task contingency slowed responses. There were no significant differences between the cohorts for either maximum d-prime (Fig. 1g), discrimination threshold (Fig. 1h), or sensitivity (all statistics can be found in Supplementary Table 1). Reaction time, however, did vary across conditions (Fig. 1i). MST mice showed shorter auditory than visual reaction times and visual reaction times decreased with increasing levels of orientation change for both UST and MST. For the same visual stimuli, reaction time was significantly longer for MST than for UST mice. For both vision and audition, reaction time negatively correlated with performance (Ext. Data Fig. 1f,g). The addition of auditory task relevance thus increases reaction times for the same visual stimuli. This result was expected because MST mice were trained to make binary decisions on whether auditory versus visual changes took place, which requires comparisons across sensory channels^36^.

### Early and late activity emerges in V1 of trained mice

To investigate whether delayed reaction times corresponded with slower dynamics of late V1 activity, we performed laminar recordings and sampled single-unit activity across cohorts (Ext. Data Fig. 2a). In NE animals the instantaneous orientation change evoked a short transient activity in V1 (until ±200 ms after stimulus onset) with a short-lasting tail (Fig. 2a; Ext. Data Fig. 2b). In visually trained animals (UST and MST), a similar transient wave occurred, but now also a late, second wave of activity was present (emerging ~200 ms after stimulus onset), primarily in hits and to a lesser extent in false alarms (Fig. 2b-f). These dynamics of early and late wave activity were seen in both threshold-level and maximal orientation changes, and in both narrow and broad-spiking cell types (Ext. Data Fig. 2).

**Figure 2:**
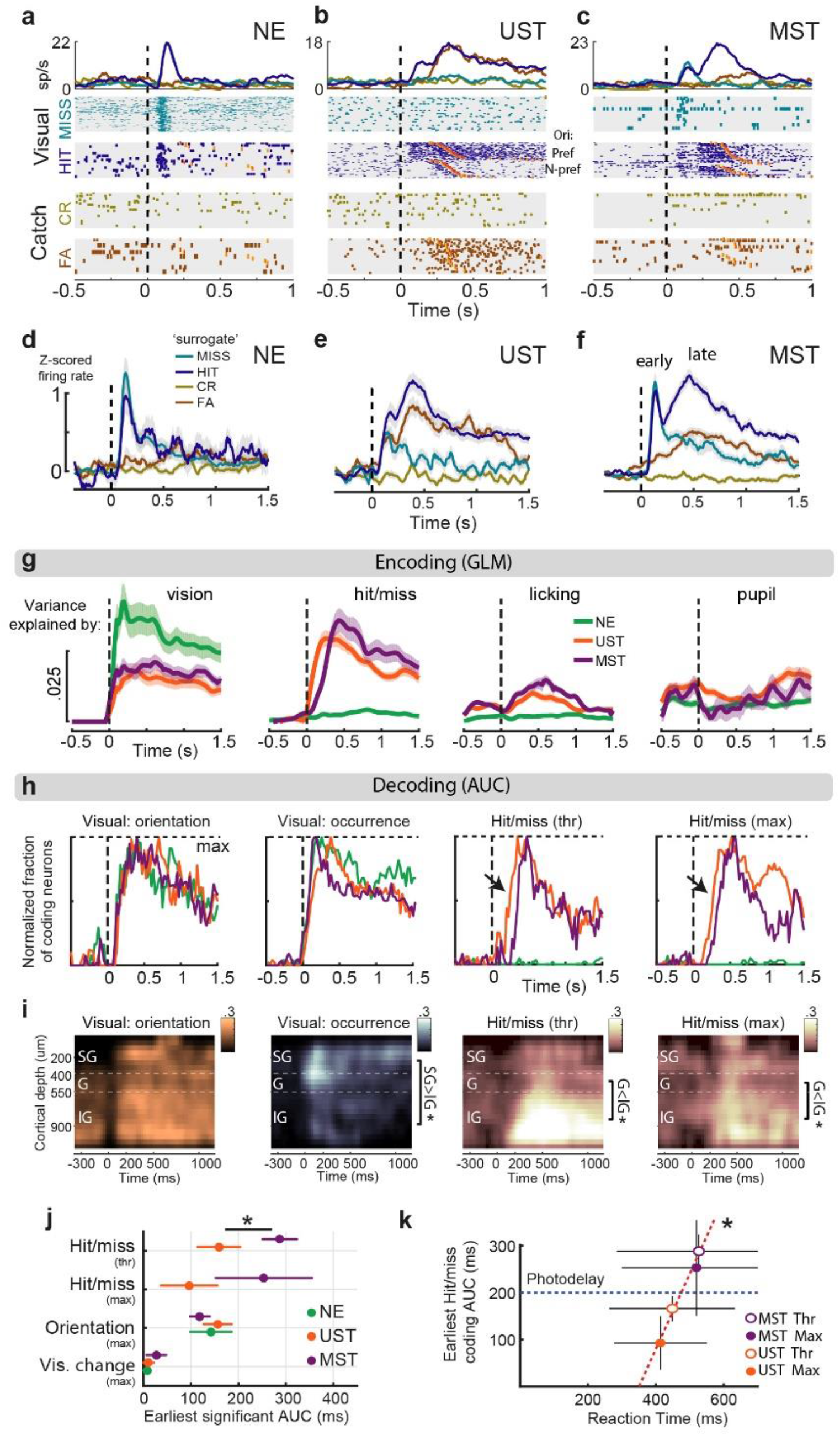
Task complexity modulates late activity in V1. a-c) Stimulus-evoked and report-related activity in three example neurons recorded in V1 (Task A). All three examples display early sensory-driven activity in visual, but not catch trials (Hit and Miss vs. CR and FA trials). Increased activity was present during the late phase (±200 to 1000 ms) in visual hit trials (c) or both hits and false alarms (b), based on individual neurons. Raster plots are grouped by trial type (visual or catch) and choice. Within trial type, trials are sorted by post-change orientation and response latency. Orange ticks show first lick and reward delivery. The top panel shows the firing rate averaged for each of these trial-type x choice combinations. The dotted line indicates stimulus change. CR = correct rejection. FA = visual false alarm. d-f) Averaging z-scored firing rate over all neurons for visual and catch trials split by choice reveals biphasic activity in visual hits but not misses, with late activity only present in animals for which visual trials were rewarded (UST and MST). Behavioral responses in NE mice should be regarded as surrogates, as visual hits in NE mice were not rewarded (see Methods). Note the increase in firing rates in FA trials for UST and MST mice but not NE mice (see Ext. Data Fig. 3 for an in-depth analysis of the lick-related nature of these responses). Line and shading indicate mean ± s.e.m. g) We constructed an encoding model in which sensory, hit/miss, movement, and arousal variables simultaneously predict firing rates of individual V1 neurons (see Ext. Data Fig. 4). Each line shows how much firing rate variance is explained for each time bin across trials based on only including a subset of predictors. In the NE mice, visual predictors (orientation and amount of change) explain a higher fraction of EV than in UST and MST mice, while movement (licks left and right) and hit/miss encoding (visual trial and correct lick interaction) make a negligible contribution. For UST and MST mice both movement and hit/miss encoding explain variance at a later onset than visual variables, and visual information remains an important predictor throughout the trial in all task versions (Statistics in Ext. Data Fig. 4). Arousal (measured in terms of pupil size) is a generally weak predictor across task versions. h) We used a single-neuron decoding analysis (ROC) to compute the fraction of neurons (summed over all recordings) coding for task-relevant variables over time. The coding of change occurrence and orientation (visual features) showed similar dynamics across cohorts, whereas hit/miss coding (visual hits vs misses) was only present in UST and MST mice (as expected) and started earlier in UST than MST mice (highlighted with black arrows). Each coding fraction is baseline-subtracted and normalized by its maximum. i) To investigate spatiotemporal coding dynamics we created heatmaps of the fraction of coding neurons across cortical depth and time. We binned neurons based on their recorded depth relative to the granular layer (400-550 μm from dura, SG=supragranular, G=granular, IG=infragranular). Orientation coding was present across cortical layers, whereas the coding of visual change occurrence was confined to an early transient wave in granular and supragranular layers (0-200 ms, SG versus IG, p=0.018). In contrast, hit/miss coding during late activity was predominant in infragranular layers relative to the granular layer (Threshold, 200-1000 ms, G vs IG, p=0.039; maximum, G vs IG, p=0.027). Only UST and MST cohorts were included to compare sensory and hit/miss coding in the same datasets. Significance (sidebars): * p < 0.05. j) The earliest increase in the fraction of significantly coding neurons was similar across cohorts for visual change occurrence or orientation, whereas hit/miss coding appeared later in MST than UST mice for threshold orientation changes (*p<0.05; maximal changes, n.s.). k) Reaction time correlated with the earliest moment of significant hit/miss coding (r=0.989, p=0.01). Each dot is the average of one visually trained condition. Red dotted line shows a linear regression fit. Blue dotted line at 200 ms marks the time point where late photostimulation was applied (see Fig. 3d). At this point, unisensory trained mice already showed hit/miss-coding in V1, while multisensory trained mice did not.

### Neural coding during late V1 activity

Recent studies have shown that late activity in V1 can reflect movement-related variables^20,21,23^. We aligned population activity to the first lick after stimulus onset and found that spiking activity across many neurons was indeed modulated by licking movements, specifically in UST and MST mice (Ext. Data Fig. 3). The amplitude of this modulation was higher in trials with correct versus incorrect licks. To further disentangle the contribution of stimulus variables (visual and auditory features and amount of change), movement variables (the timing and number of lick responses), the hit/miss distinction (hit, with reward, or miss, without reward), and arousal (pupil size), we built a kernel-based generalized linear model (GLM)^22,37,38^ where we included these variables as predictors of firing rate (Ext. Data Fig. 4). We computed the cross-validated variance of firing rate explained over time by each of these subsets of predictors (Fig. 2g; Ext. Data Fig. 4e-g). In NE mice, visual predictors explained most of the variance. In UST and MST mice, besides visual predictors, we found that, during the 200-1000 ms post-stimulus window, hit/miss and movement both explained a significant portion of the variability. In sum, late V1 activity reflected a combination of visual, movement, and hit/miss-related variables, but only in trained mice.

**Figure 3:**
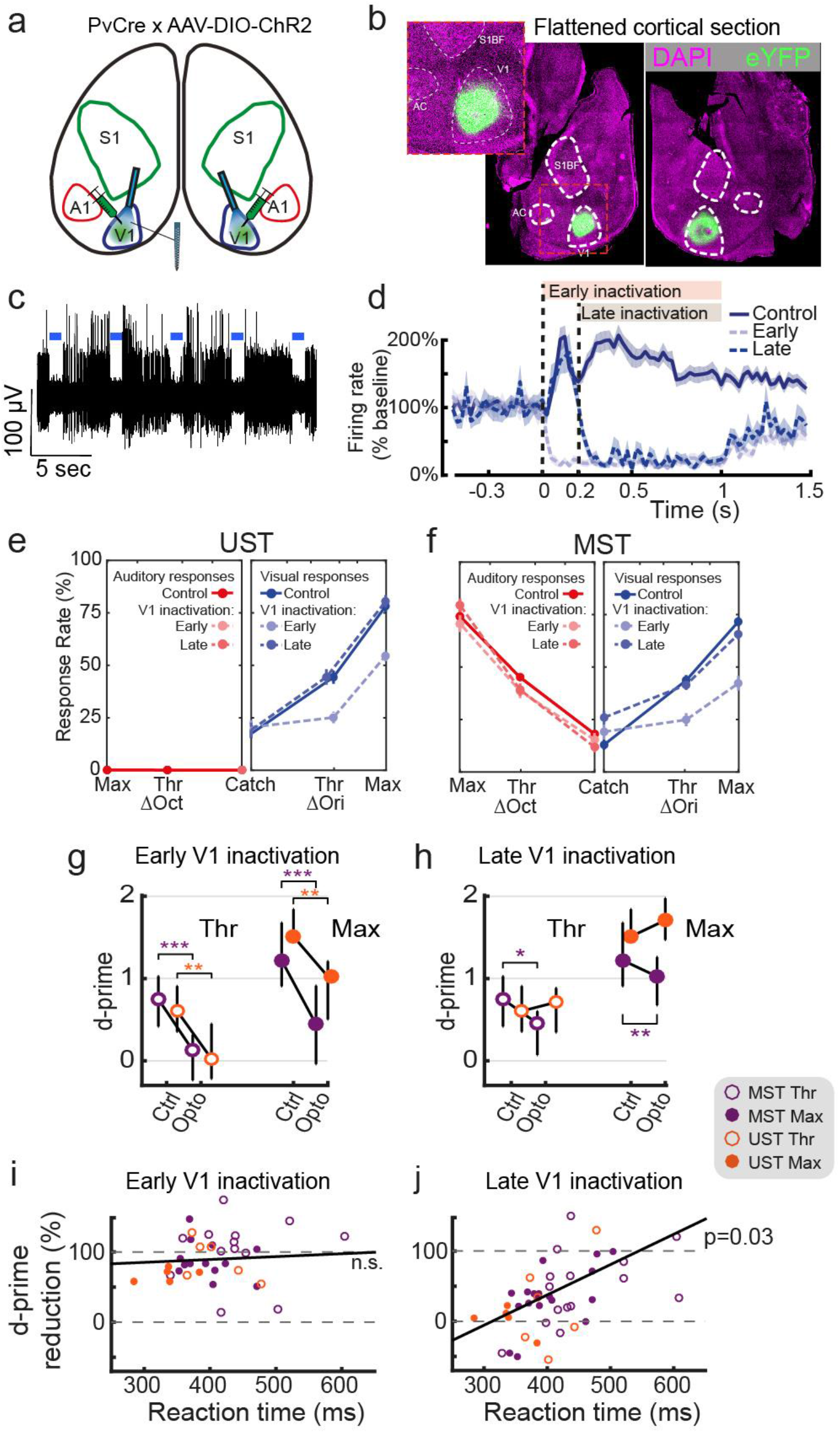
Late silencing of V1 selectively impairs task performance of sessions with slow reaction time. a) Schematic of optogenetic strategy in Task A. Cre-dependent ChR2 expression in bilateral V1 of PvCre mice allowed robust silencing by locally enhancing PV-mediated inhibition. A1 = auditory cortex, S1 = primary somatosensory cortex. b) Dorsal view of flattened cortical hemispheres sectioned approximately through layer 4 showing localized viral expression in bilateral V1. c) High-pass filtered trace from an example V1 recording site showing robust silencing of multi-unit spiking activity during bouts of 1-second photostimulation (blue bars). d) Photostimulation silences V1 in a temporally specific manner. Early blue light stimulation (i.e. starting at the onset of stimulus change) reduced the activity of putative excitatory neurons (see Ext. Data Fig. 2f,g) to about 5% of their baseline activity. Late photostimulation left the initial sensory-driven response intact but silenced activity after 200 ms relative to stimulus onset. Traces show baseline-normalized firing rate averaged over V1 neurons from UST and MST mice. Control trials are visual hits. e) Early, but not late, silencing impaired threshold (Thr) and maximal (Max) visual change detection in UST mice. Behavioral detection rates for control, early, and late silencing trials follow plotting conventions of Fig. 1d-f. f) Both early and late silencing affected visual change detection rates in MST mice. For the increase in FA see Methods. g) Early silencing affected visual discrimination performance (d-prime) for both saliencies across UST and MST cohorts. *p<0.05, **p<0.01,***p<0.001. h) Effect of late silencing depended on task type: late silencing only reduced d-prime in MST, but not UST mice. i) The effect of early silencing (quantified as the reduction in d-prime) was not significantly correlated with the median reaction time in control trials from the same session (r=0.048, p=0.865). j) Same as *i* but for late silencing. The effect of late silencing was significantly correlated with the reaction time (r= 0.423 p=0.03).

**Figure 4:**
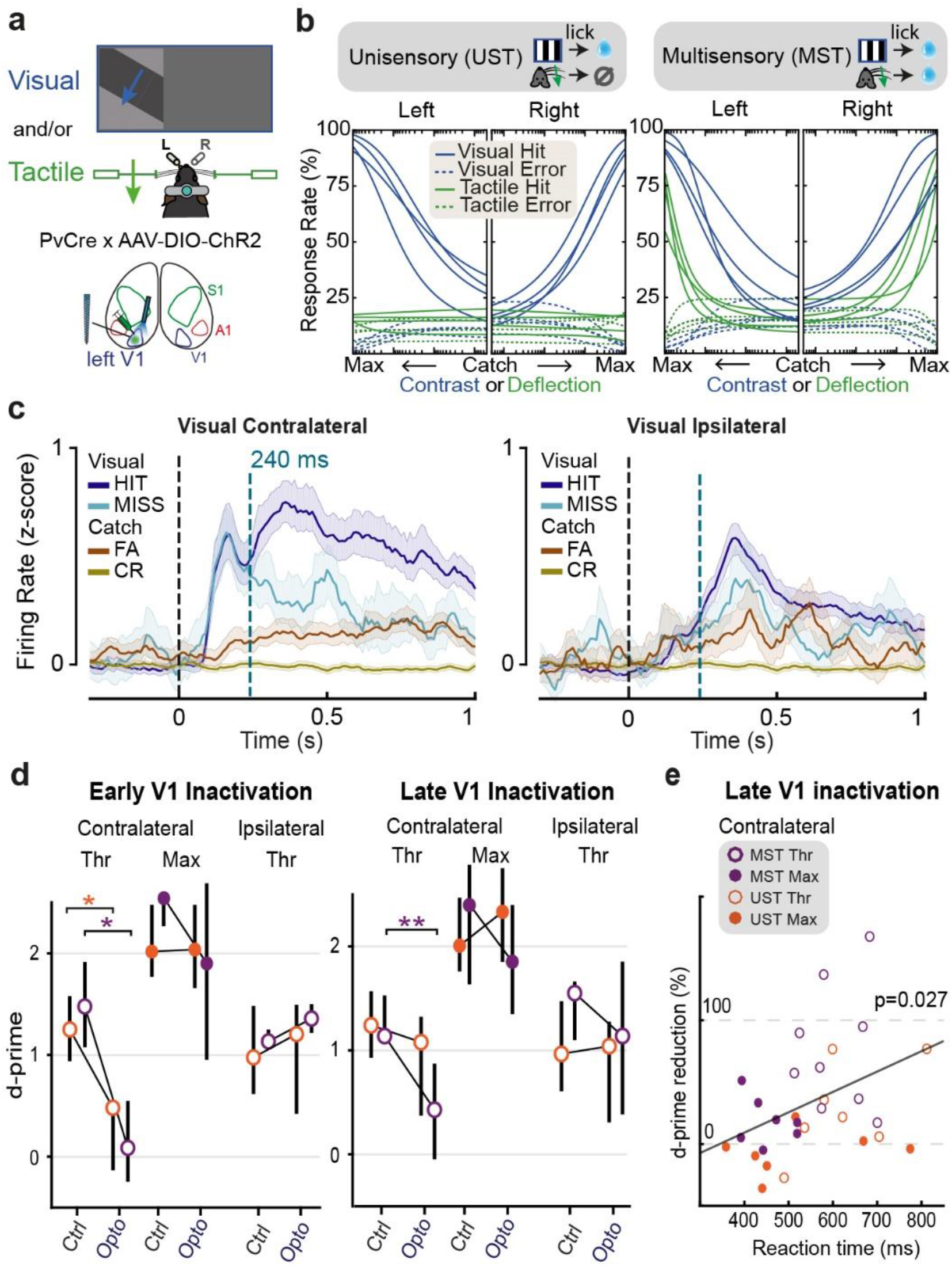
Extended causal requirement of V1 generalizes to visuotactile side detection. **a)** Scheme of the visuotactile two-sided detection task in which mice reported the side of visual and/or tactile stimulation. In this task B, visual and tactile information need to be integrated as an inclusive- or operation (rather than discriminated as in task A) to decide as to which side (left/right) was stimulated. **b)** Psychometric fits for visual and tactile detection for each mouse trained in the UST and MST version of the task. The horizontal axis represents stimulus intensity. The origin lies in the center, representing catch trials. The axes run to the left for stimuli presented to the left and right for stimuli presented to the right. Intensity values represent contrast for visual trials and deflection angle for tactile trials. They were normalized for rendering purposes. **c)** Average z-scored firing rate of responsive V1 neurons during visual-only and catch trials, split by trial outcome. Contralateral stimuli elicit biphasic activity in hits but reduced late activity in misses. The traces shown correspond to maximum-saliency stimuli and MST mice. The onset of optogenetic silencing at 240 ms is indicated by a blue dotted line. Shaded area: bootstrapped 95% confidence intervals. **d)** Median task performance (d-prime) during early (left) and late (right) V1 silencing for different visual stimulation conditions. Thr: threshold-level stimulus saliency; Max: maximum stimulus saliency. V1 silencing had no effect on performance for ipsilateral threshold-level stimuli nor contralateral maximum-level stimuli. It only affected the detection of contralateral threshold-level stimuli. While early silencing impaired visual detection performance in both UST and MST (p=0.031 for both), late silencing only affected MST mice (p=0.008). For ipsilateral threshold and contralateral maximal stimulation, early or late silencing V1 had no effect, only for contralateral threshold stimulation. Error bars: inter-quartile interval. **e)** Reduction in d-prime by late silencing correlated with the median reaction time on corresponding control trials, per session (p=0.027, r^2^=0.162).

### Multisensory context delays the time course of late activity

To identify how the delayed reaction time in MST mice was associated with coding dynamics of single neurons we used a receiver operating characteristic (ROC) analysis^39,40^. Across task versions, the ratio of neurons coding for visual features (grating orientation, and occurrence of a visual stimulus change – i.e. visual trials versus trials with no stimulus change, catch trials) was similar across cohorts. In UST and MST mice, however, visual report (i.e. hits vs. misses) was also encoded by a substantial fraction of neurons, in line with our regression model (Ext. Data Fig. 5a). To understand at which time points visual features and hit/miss coding could be read out, we plotted the fraction of neurons that significantly coded for each of these variables over time (Fig. 2h; Ext. Data Fig. 5b). Temporal dynamics were strikingly similar across cohorts for sensory variables, while hit/miss coding appeared later in V1 for MST than UST mice. Sensory and hit/miss coding were spatially segregated across cortical depths, suggesting that these two processes have different neural substrates. Coding of visual change was predominant in superficial layers, and hit/miss coding in deeper layers (Fig. 2i; Ext. Data Fig. 2e). We quantified the earliest moment of a significant increase in the fraction of coding neurons relative to baseline and found that only hit/miss-coding was delayed in MST compared to UST (Fig. 2j; threshold changes: 288 ms ± 36 ms versus 162 ± 28 ms, MST vs. UST, p<0.05; maximal changes: 249 ± 104 ms vs 92 ± 56 ms, MST vs. UST, n.s.). In relation to the delayed visual reaction times in MST mice, we found a strong correlation between the onset of hit/miss-coding and reaction time. Hit/miss-related activity preceded the first lick by ±280 ms (Fig. 2k, Methods). Therefore, at 200 ms after stimulus onset (blue dotted line in Fig. 2k) UST mice already showed hit/miss-coding in V1, while MST mice did not.

**Figure 5:**
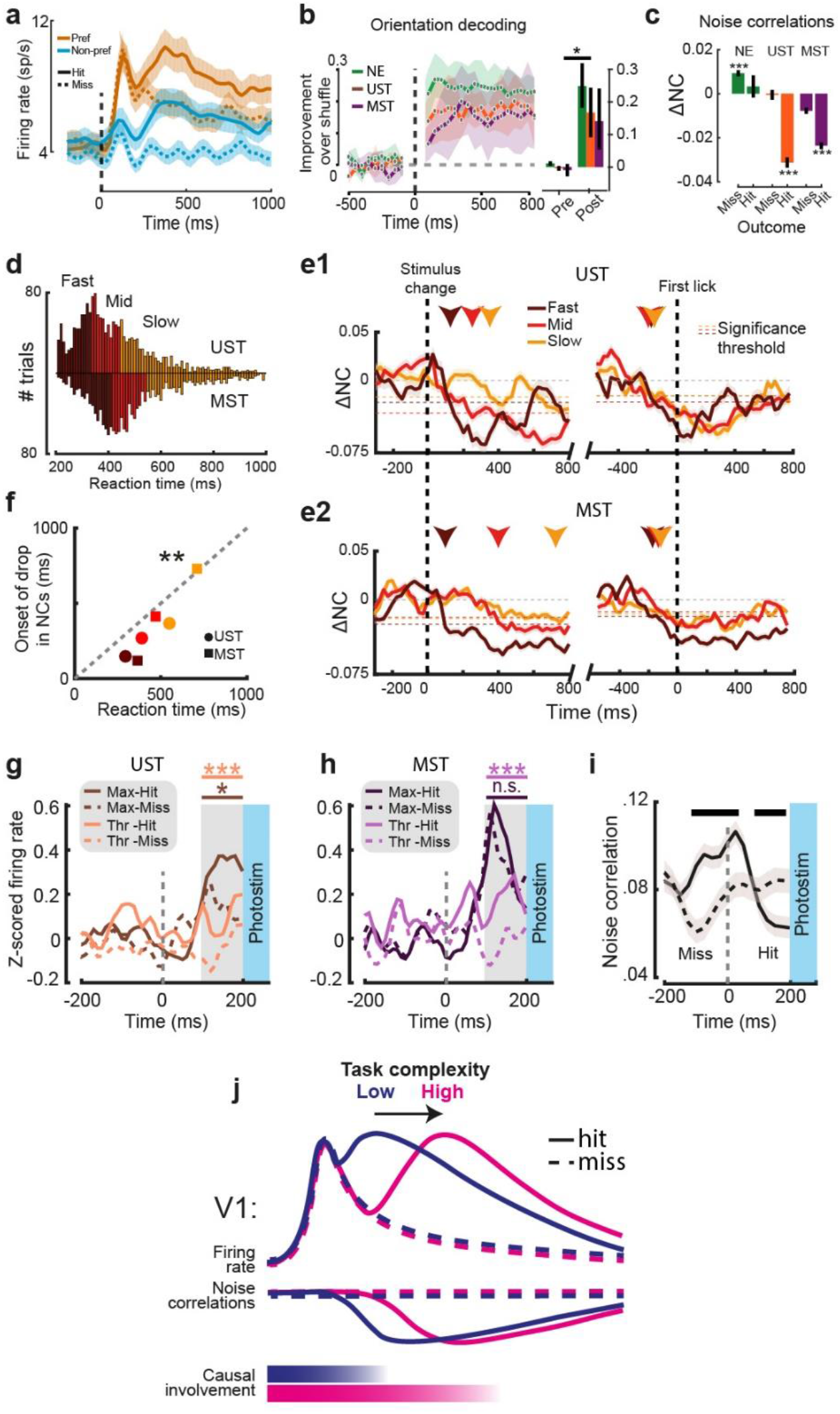
Drop in noise correlations predicts effects of late silencing. a) Average spiking rate for all orientation-selective neurons for preferred and non-preferred orientations split by hits and misses (UST and MST neurons combined; task A). Note how orientation-selective neurons show a late hit-modulation of firing rate. b) We trained decoders to discriminate post-change grating orientation (all visual trials) from V1 population activity. Decoding performance increased after stimulus change similarly in all cohorts. Inset shows increased decoding performance for post-stim (0 to +500 ms) versus pre-stim (−500 to 0 ms; p=0.00195, cohorts combined). c) Change in noise correlation (NC) relative to baseline (200 to 1000ms compared to baseline −500 to 0 ms) for visual trials split by choice and cohort (for auditory trials, see Ext. Data Fig. 10b). Noise correlations decreased only during hits in UST and MST mice (p<1e-17, p<1e-18, respectively). Misses in NE mice were associated with a slight increase in noise correlations (p<1e-4). d) Visual hit trials were separated into three tertiles based on reaction times to investigate reaction time-dependent noise correlation dynamics. The histograms show the reaction time distributions for visual hits in UST and MST cohorts and tertile ranges. e1) The onset of the drop in NCs in UST mice was related to reaction time. In the left panels, noise correlations (aligned to stimulus change) drop – relative to baseline (below 2 standard deviations of the baseline; −500 to 0 ms) – first in fast trials, and progressively later in medium and slow trials. Right panels show that, when aligning to lick onset, the drop in noise correlations precedes reaction times by a similar lag, independent of reaction time tertile. Horizontal dashed lines indicate, for each tertile, the threshold used to determine the onset of the drop in NCs, highlighted with colored arrows. e2) Same as e1, but for MST mice. f) Reaction time and moment of decorrelation were significantly correlated (r=0.960, p=0.002). Scatterplot shows median reaction time and earliest time point of decorrelation for each tertile in the two visually trained cohorts. g) Firing rates just before photostimulation (100-200 ms) were higher if the trial was a visual hit rather than a miss in UST mice (Thr: p=0.001; Max: p=0.029). Plot shows average Z-scored rate of both putative excitatory and inhibitory neurons during visual trials with late photostimulation, split by hit/miss. h) Same as g, but for MST mice. Firing rates just before photostimulation were higher for hits than misses only for threshold visual changes (p=0.001), but not maximal changes (p=0.34). i) Noise correlations for visual hit and miss trials before photostimulation onset (grouped across UST and MST cohorts and saliency levels). Black bar on top: time bins with significantly different noise correlation values between hits and misses (p < 0.05). Throughout the figure, lines and shading are mean ± SEM. j) Schematic summary of results: increased task complexity (such as through multisensory interactions) delays the onset of the late report-related wave of activity and drop in noise correlations and extends the causal involvement of V1. Jointly, these processes may enhance the effectiveness of V1 in transferring task-relevant information to downstream areas.

### Late activity is causally required for perceptual decision making

Next, we wondered whether V1 activity occurring after the onset of report-related activity could be causally linked to perception. We locally expressed ChR2 in parvalbumin-expressing interneurons in V1 (Fig. 3a,b, Ext. Data Fig. 6a) to achieve temporally specific optogenetic inactivation^41,42^. Laser stimulation robustly silenced multiunit activity (Fig. 3c). To determine the temporal window of V1 involvement, we silenced it either from the moment of stimulus change onwards (*Early*, 0 ms) or from the 200 ms temporal cutoff we identified in the onset of hit/miss coding in UST and MST mice (*Late*, 200 ms; Fig. 2k). Pyramidal cell activity in V1 was silenced to around 5% of spontaneous baseline activity in a temporally specific fashion (Fig. 3d).

Early silencing of V1 strongly reduced detection of orientation changes during both UST and MST task performance (Fig. 3e-g), consistent with the primary role of V1 in visual feature processing^26,43,44^. Interestingly, late silencing only affected change detection performance of MST mice (Fig. 3h). V1 silencing did not affect auditory performance (Ext. Data Fig. 6b). Moreover, photoillumination of control area S1 did not affect visual or auditory performance (Ext. Data Fig. 7).

Even though late silencing impaired visual detection in MST mice *on average*, results across animals and experimental sessions were mixed: some sessions showed robust behavioral impairment, whereas others showed little effect (Ext. Data Fig. 6c). Thus, we investigated whether variability in reaction speed – with reaction speed being a proxy for subjective task complexity (Ext. Data Fig. 1f,g) – could underlie the divergence in the effectiveness of late silencing. We plotted the reduction in d-prime as a function of reaction time (Fig. 3i,j; Ext. Data Fig. 6d,e). Whereas early silencing invariably reduced performance, the effect of late silencing scaled with reaction time (Fig. 3j). This was not the case for the animal’s propensity to lick (affecting false alarms and quantified with the criterion parameter in our behavioral model, Ext. Data Fig. 6f), suggesting that perceptual sensitivity was specifically affected in slow, subjectively more complex, experimental sessions. Late silencing thus left performance intact in ‘fast’ (and subjectively easier) sessions in which hit/miss coding emerges quickly (mostly UST sessions) and reduced performance in ‘slow’ sessions where hit/miss coding started after 200 ms (mostly MST sessions, but note also how one slow UST session was affected – Fig. 3j).

### Causal involvement of late activity generalizes to visuotactile side detection

So far, our results suggest that in the multisensory variant of the change detection task (MST), late V1 activity is causally involved whereas in the unisensory variant it mostly is not. However, UST and MST cohorts do not only differ by sensory contingencies, as UST mice were trained on a Go/No-Go paradigm, while MST mice learned a two-alternative choice task. Thus, the results we report could be due to differences in behavioral strategy rather than to changes in multisensory context. Furthermore, we wondered whether our results may extend to other sensory modalities. To address these aspects, we developed a visuotactile side detection task in which mice reported the side of sensory stimulation, i.e. instructing them to lick left for visual or tactile stimuli presented to the left and oppositely for the right side (Task B – Fig. 4a). Stimuli consisted of monocular drifting gratings (visual), whisker pad deflection (tactile), or a combination of both. Again, some mice were trained on responding only to vision to obtain reward (UST), while another cohort was trained on both vision and somatosensation (MST).

Importantly, this UST version still contained two response options and required responding to the correct lick spout (the visual stimulus could appear on the left or right). In addition to the differences with task A, this new task allowed us to test if our results extended to another multisensory processing principle (congruent combination of modalities instead of segregation^29^) and the detection of a different stimulus dimension (contrast instead of orientation change).

We controlled stimulus salience by varying visual contrast and whisker deflection amplitude and fitted the behavioral data with a psychometric model (Fig. 4b, see Methods). The visual detection threshold and task performance at maximum saliency were similar for both cohorts (UST and MST, Ext. Data Fig. 8a-b). As in Task A, we found that visual reaction time (RT) decreased for higher stimulus saliencies (Ext. Data Fig. 8c). In contrast with task A, however, RTs were similar between tactile and visual trials.

Pursuing the comparison with task A, laminar recordings in V1 revealed similar neural dynamics, with a marked early stimulus-driven component visible in contralateral visual trials, and late activity in both contra- and ipsilateral visual hits (Fig. 4c, Ext. Data Fig. 9a-c). This late activity was also present in tactile hits, although only for MST mice (Ext. Data Fig. 9d). We optogenetically silenced unilateral V1 either from stimulus onset (early silencing) or after a delay that separated the late from the early wave of activity (240 ms, late silencing; Fig. 4c). Early silencing of unilateral V1 reduced the detection of contralateral threshold-contrast stimuli in both UST and MST mice but spared detection of ipsilateral stimuli and full contrast stimuli (Fig. 4d). In MST mice, tactile detection was not affected by V1 silencing (Ext. Data Fig. 8d). Consistently with our results for Task A, while early V1 silencing impaired detection of threshold-level visual stimuli for both unisensory and multisensory contexts, late V1 silencing only affected such detection in MST mice (Fig. 4d). As in Task A, we observed that the effect of silencing increased for conditions with longer reaction time (Fig. 4e). Overall, results obtained with task B generalize our findings and confirm that the temporal window for the causal involvement of V1 in perceptual decision making is extended when subjects reinstate the more complex, multisensory attentional set they have been trained on.

### Population decorrelation predicts late silencing effects

The late report-related wave of activity during visual hits (Figs. 2, 4d) likely arises through an interplay of higher-order areas that feed back to V1, possibly including premotor or other frontal areas^18,20,23,45^. The timing of this wave predicts the behavioral effects of late V1 inactivation, but the underlying mechanism governing the sculpting of a behavioral decision remains unclear. One possibility is that late activity is predominantly related to movement variables, coded orthogonally to sensory representations from a population perspective^21^. To investigate this, we further explored the properties of late activity. First, we tested whether the extended causal requirement of V1 was related to changes in the fidelity of sensory processing, as indexed by orientation decoding. For the audiovisual change detection task, we examined the effect of report-related activity modulation on orientation-tuned neurons (Fig. 5a) at a population level and trained a random forest decoder to decode post-change grating orientation from V1 population activity. Orientation decoding was possible for hundreds of milliseconds after the orientation change with comparable performance across the three task versions (Fig. 5b). This suggests that in all visual trial types (regardless of task contingencies) information regarding the orientation of the stimulus was similarly present and that the extended requirement of V1 could rather be due to the interaction of this representation with the rest of the cortical circuit.

Correlated firing rate fluctuations that are unrelated to signal coding (noise correlations – NCs) can impact information coding in populations of neurons^46–48^. NCs decrease as a function of various conditions, for instance when animals become experts at change detection^49^ or through attention^50^. Such changes in noise correlations may allow V1 representations to more effectively drive downstream targets^51^. We computed pairwise firing rate fluctuations over time for visual hits and misses. During baseline (−500 to 0 ms) NC values were comparable to the literature^52–55^ (0.063 ± 0.14 std) and decreased after stimulus change only during hits in UST and MST but not NE mice (Fig. 5c, Ext. Data Fig. 10a). To investigate whether the onset of the decorrelation was related to reaction time, we split all visual hits from V1 recording sessions into three tertiles based on reaction time (Fig. 5d). Similar to behavioral data without recordings (Fig. 1i), UST mice reacted faster than MST mice (p=1.8e-68). We quantified the earliest time-point where the drop in NCs reached significance (relative to baseline) for each tertile for UST and MST mice. NCs decreased at a latency that depended on reaction time, with the drop in NCs occurring later on slow compared to fast trials (Fig. 5e). The latency of the decrease in NCs and reaction time were significantly correlated (Fig. 5f), suggesting that population decorrelation is time-locked to reaction time. Indeed, noise correlations relative to the first lick (see Methods) dropped just preceding this first lick, irrespective of reaction time (Fig. 5e, right part; Ext. Data Fig. 10b), with the strongest decrease for visual hits in UST and MST mice and no decrease in surrogate hits in NE mice (Ext. Data Fig. 10c). Overall, this suggests that a late drop in NCs is not simply a correlate of (preparatory) motor activity, but may rather play a mechanistic role in perceptual report.

### Activity level and decorrelation predict the effect of late silencing

If the late-onset drop in NCs contributes to perceptual report, one would expect that the variability in the behavioral effect of silencing (i.e. whether V1 inactivation was followed by a hit or miss) could be explained at the single-trial level by whether this drop had already occurred at the time of photostimulation. We therefore focused on V1 activity during visual trials just before late inactivation started (n=230 cells), and tested whether the report-related firing-rate modulation and drop in NCs had both already occurred before 200 ms in hits but not misses. Indeed, hits were associated with increased activity just before photostimulation started (100-200 ms after stimulus onset) across levels of stimulus change and task versions (Fig. 5g-h). Similarly, NCs showed distinct profiles for hits and misses (Fig. 5i) and decreased just before silencing onset during hits but not misses. Surprisingly, NCs were higher just before and after stimulus change for hits (−125 to +25 ms around stimulus change), in contrast with^56^. Overall, these results show that increased firing rates, and the temporally coinciding drop of NCs in V1, may both contribute to an improved readout and communication of visual activity to downstream areas (Fig. 5j).

## Discussion

In this study, we investigated the nature and function of late, recurrent activity in V1 in perceptual decision-making. An increase in subjective task complexity delayed behavioral decisions and extended the temporal window in which V1 was causally involved in determining perceptual report, given the same visual stimulus. As animals in the MST tasks were trained to process behaviorally relevant signals in two sensory modalities rather than one, longer reaction times (compared to UST tasks) are likely needed to integrate and compare information from distinct sensory modalities, also to assess which of two modalities is most likely to present an externally (as opposed to self-) induced sensory change. Similarly, a specific visual stimulus under conditions of low saliency (i.e., a small change in grating orientation) likely requires more time to determine whether there was a change. Therefore, our results may not be specific to multisensory contexts (cf. Fig. 3j), as any condition increasing cognitive demands might delay the contribution of dorsal cortical areas to decision making, including an increased reliance on V1^57^.

We found modest differences in onset latencies and orientation coding of visually evoked V1 responses across task contingencies and cohorts, suggesting that the dynamics of bottom-up, feedforward processing are mostly conserved. In contrast, striking differences were found in the late phase of V1 activity, and particularly in the behavioral consequence of late optogenetic inactivation. Late activity is thought to arise from recurrent feedback emitted by higher-order cortices^32^, in agreement with the predominance of report-related coding in deeper cortical layers (Fig. 2i). Silencing late V1 activity abolished detection of orientation changes and contralateral stimuli in conditions of high task complexity (and consequently long reaction time). This effectiveness of late inactivation was most prominent for low-saliency stimuli in MST mice and linearly scaled with reaction times (Fig. 3j, 4g). The only exception was the detection of high contrast stimuli in MST (but also UST) mice in task B, which was affected neither by early nor late V1 silencing (Fig. 4d,e). This suggests that subcortical structures (e.g. superior colliculus) may suffice to localize highly salient stimuli^58^ in task B (in contrast to task A, which requires detecting an orientation change), although we cannot fully exclude that portions of V1 were not completely inactivated. Furthermore, it is unlikely that late V1 silencing generally impaired cortical network processing^59^, as it did not affect ipsilateral visual detection in the visuotactile detection task (task B; Fig. 4), nor tactile or auditory performance (Fig. 3 and Ext. Data Fig. 4d).

We optogenetically silenced both sensory and report-related components of V1 activity, which are jointly present in the late window (Fig. 2g). Importantly, late V1 inactivation in the absence of an early sensory-related component (e.g. ipsilateral visual stimuli in task B, or non-visual hit trials) did not impair behavioral responses, in agreement with a recent study suggesting that the report-related component alone is not sufficient for perceptual task performance^17,44^. However, the question remains how sensory-evoked and report-related activity are related to each other during the late phase^44^. On the one hand, higher task complexity may prolong sensory processing of the visual stimulus, or at least the time downstream regions need to sample the ongoing flow of V1 activity to gather sufficient evidence on visual stimulus change^60,61^. On the other hand, recurrent interactions between visual cortex and connected regions during late windows may jointly influence sensory representation, in line with the predictive processing framework^15,32,62^. In other words, in subjectively complex task conditions V1 and downstream regions need to interact for longer periods to jointly construct a behaviorally conclusive representation of the modality-specific change (task A) or the side of stimulus presentation (task B). Notably, we increased task complexity but kept visual stimuli unchanged. Therefore, if V1 was merely passively transmitting information to higher-order areas, it is unlikely that we would have observed a task-related change in activity mode, represented by a co-occurring increase in firing rates and drop in NCs. Conversely, we found that the increase in the amplitude of late activity and drop in NCs both preceded behavioral reactions by a relatively constant lag (Fig. 2k, 5f), and predicted the effect of late optogenetic inactivation (Fig. 5g-i). This suggests that, rather than being simply sampled for longer periods by downstream regions, V1 activity is actively modulated as a function of task complexity. Together, the temporal relationship between the onset of the hit-miss modulation of firing rate and the decrease in NCs (both preceding RT during hit trials, cf. Fig. 2k and Fig. 5f), and the trial-by-trial efficacy of late silencing, point towards a link between the two processes (Fig. 5j), which however remains to be causally investigated. The increased population activity (Fig. 2) and the co-occurring drop in NCs (Fig. 5), may jointly play a role in shaping information flow across the interconnected cortical network ranging from V1 to frontal regions^45,62,63^. The function of both processes may therefore be to enhance the effectiveness of V1 in transferring task-relevant information to downstream areas^64^.

In conclusion, our results show that, although all sensory information that is theoretically required to perform a task is available in V1 shortly after stimulus onset, transforming such sensory inputs into a perceptual representation requires substantial recurrent interplay between cortical areas, which is temporally extended by factors increasing task complexity (such as multisensory interactions). Late-onset activity in primary visual cortex is therefore not simply involved in relaying, refining, and modulating the processing of complex visual stimuli, but also provides a crucial contribution to perceptual decision-making.

Our results thus dispute the classical picture of perceptual decision making: V1 is not only needed to relay processed visual information to higher-order areas, but continuous interactions between V1 and downstream regions are required to solve complex perceptual tasks.

## Acknowledgments

We thank D. Sridharan for providing code for the multi-alternative detection model; C. Rossant, members of the Cortex Lab (UCL) and contributors for Klusta and Phy spike sorting software; Andriana Mantzafou, Klara Gawor, and Alexis Cervàn Canton for assistance in behavioral training. This work was supported by the European Union’s Horizon 2020 Framework Program for Research and Innovation under the Specific Grant Agreement 720270 (Human Brain Project SGA1) to C.M.A.P., Grant Agreement 785907 (Human Brain Project SGA2) and 945539 (Human Brain Project SGA3) to C.M.A.P. and U.O., by the FLAG-ERA JTC 2015 project CANON (co-financed by the Netherlands Organization for Scientific Research – NWO) to U.O., by the FLAG-ERA JTC 2019 project DOMINO (co-financed by NWO) to U.O. and by the Amsterdam Brain and Mind Project to C.M.A.P. and C.P.K.

## Author contributions

Conceptualization, M.O.L., J.P., C.M.A.P., U.O.; Methodology, M.O.L. and J.P.; Main analysis, M.O.L. and J.P., Additional analysis, P.M., J.M.; Data Curation, M.O.L. and J.P.; Writing – Original Draft, M.O.L., J.P, U.O.; Writing – Review & Editing, All; Visualization, M.O.L. and J.P.; Supervision, U.O., C.M.A., and C.P.K.; Funding Acquisition, U.O., C.M.A. and C.P.K.

## Methods

### Data and Code Availability

The data and code that support the findings of this study are available from the corresponding author, U.O., upon reasonable request.

### Animals

All animal experiments were performed according to national and institutional regulations. The experimental protocol was approved by the Dutch Commission for Animal Experiments and by the Animal Welfare Body of the University of Amsterdam. We used two transgenic mouse lines: PVcre (B6;129P2-Pvalbtm1(cre)Arbr/J, JAX mouse number 008069) and PVcre/TdTomato (C57BL/6-Tg(Pvalb-tdTomato)15Gfng/J, JAX mouse number 027395). A total of 49 male mice (28 in task A, 12 in task B) were used for this study. Mice were at least 8 weeks of age at the start of experiments. Mice were group-housed under a reversed day-night schedule (lights were switched off at 8:00 and back on at 20:00). All experimental procedures were performed during the dark period. This study did not involve randomization or blinding. We did not predetermine the sample size. Some subjects were unable to successfully learn to make decisions based on both modalities (MST task versions) within 2 months and were excluded from further experiments.

### Head-bar surgery

Before the start of any experiments, mice were implanted with a headbar to allow head-fixation. Mice were subcutaneously injected with the analgesic buprenorphine (0.025 mg/kg) and maintained under isoflurane anesthesia (induction at 3%, maintenance at 1.5–2%) during surgery. The skull was exposed and one of two types of custom-made titanium circular head-bars with a recording chamber (version 1: inner diameter 5 mm, version 2: inner diameter 10 mm) was positioned over the exposed skull to include V1 and attached using C&B Super-Bond (Sun Medical, Japan) and dental cement. For task A in which visual stimuli were centrally presented binocular V1 was targeted based on the following coordinates (relative to lambda): AP 0.0, ML +/− 3.0^65^. Whereas coordinates sufficed for task A, for task B, in which lateralized visual stimuli were used, V1 was targeted using intrinsic optical imaging (see below) to localize the retinotopic region of V1 corresponding to the region of visual space in which the lateralized visual stimuli were presented. The skin surrounding the implant was covered using tissue adhesive (3M Vetbond, Maplewood, MN, United States) to prevent post-surgical infections. The recording chamber was covered with silicon elastomer (Picodent Twinsil). Mice were allowed to recover for 2-7 days after implantation, then habituated to handling and head-fixation before starting on the training procedure.

### Behavioral training

Mice were subjected to a water restriction schedule and minimum weight was kept above 85% of their average weight between P60-P90. They typically earned their daily ration of liquid by performing the behavioral task but received a supplement when the earned amount was below a minimum of 0.025 ml/g body weight per 24h.

Mice were head-fixed in a custom-built headbar holder in a dark and sound-attenuated cabinet. The body of the mouse was put in a small tube to limit body movements. The task was controlled in Octave interfacing with Arduino microcontroller boards. Licks were detected by capacitance-based or piezo-electric-based detectors. Upon correct licking, i.e. in hit trials, 5-8 μl of liquid reward (infant formula) was delivered immediately using gravitational force and solenoid pinch valves (Biochem Fluidics). Reward volume was calibrated biweekly to prevent lateralized response bias due to unequal reward size.

### Behavioral task A: Audiovisual Change Detection

#### Visual Stimuli

Visual stimuli consisted of full-field drifting square-wave gratings that were continuously presented with a 60 Hz refresh rate on an 18.5-inch monitor positioned at a straight angle with the body axis from the mouse at 21 cm from the eyes. Gratings were presented with a temporal frequency of 1.5 Hz and spatial frequency of 0.08 cycles per degree at 70% contrast and gamma-corrected. In trials with a visual change the orientation of the drifting grating was instantaneously changed (e.g. from 90° to 120°) while preserving the phase. The degree of orientation change determined the visual saliency and was varied across experimental conditions.

#### Auditory Stimuli

The auditory stimulus was a stationary Shepard tone, which is composed of a center frequency and a series of harmonics. In our case we used 5 harmonic tones with 2 lower and 2 higher harmonics (if f_0_ is the center frequency: f_−2_=¼*f_0_, f_−1_=½*f_0_, f_+_1=2*f_0_; f_+2_=4*f_0_). The total stimulus set of center frequencies was expressed in scientific pitch and consisted of a full octave spanning from 2^13^ Hz (=8372 Hz, which corresponds to a C_9_) to 2^14^ Hz (=16744 Hz, corresponding to C_10_) in steps of 1/256 partial octaves. For each given Shepard tone in this stimulus set, the weight of the center and harmonic tones are taken from a fixed Gaussian weight distribution over all center and harmonic tones, in this case centered at 2^13.5^ (=11585 Hz). Because of this fixed weight distribution, the 2^13^ Shepard tone is equal to the 2^14^ Shepard tone (both are a compound of 2^11^, 2^12^, 2^13^, 2^14^, 2^15^, 2^16^ with the same fixed weights) and the total stimulus set (from 2^13^ to 2^14^) is therefore circular. In trials with an auditory change, the stimulus was changed instantaneously from one stationary Shepard tone to another, i.e. to a stimulus with a different center and harmonic frequencies. For example, an auditory change of ¼ octave would jump from 2^13^ to 2^13.25^ (this was as salient as 2^13.75^ to 2^13^, due to the circularity at 2^13=14^). The degree of frequency change determined the auditory saliency and was varied across experimental conditions. The phase across all center and harmonics was preserved during auditory stimulus changes. Stimuli were presented with a sampling rate of 192 kHz. Stimuli were high-pass filtered (Beyma F100, Crossover Frequency 5-7 kHz) and delivered through two bullet tweeters (300 Watt) directly below the screen. Sound pressure level was calibrated at the position of the mouse and volume was adjusted per mouse to the minimum volume that maximized performance (average ±70 dB).

In an earlier cohort of mice trained on task A (N=13/28), the Shepard tones (1) were expressed in absolute Hz (e.g. an auditory trial with Δ2kHz changed from 8 kHz to 10 kHz), (2) had 9 harmonics, (3) were presented with a sampling rate of 48 kHz and (4) were not phase-preserved during a change in auditory frequency. We observed no qualitative or quantitative differences in both neural and behavioral results between the cohorts (behavior between cohorts is compared in Ext. Data Fig. 1). The horizontal axes were normalized in Fig. 1 to accommodate all mice.

With both auditory and visual stimulus sets being circular, the direction of change (clockwise or anticlockwise) was behaviorally irrelevant (isotropy), and the only relevant dimension was the amount of change. Given the use of the full auditory spectrum and full-field visual gratings, stimuli in both modalities allowed change detection based on feature selectivity while recruiting neurons across the tonotopic organization of auditory cortex^66^ and across the retinotopic map of visual cortex x - which in our case benefitted both neural data acquisition and interventions.

#### Versions of Task A

Animals were assigned to one of three versions of a change detection task (NE, UST, MST) in which the visual and auditory stimuli were identical and only the reward contingencies varied.

##### *NE: Noncontingent exposure* (N=7/28 animals)

In this version, neither modality was associated with reward availability. Both the auditory and visual stimuli were continuously presented with the same distribution of trial types and temporal statistics as the multisensory version (see below). To compare intermittent licks, rewards, and stimuli across task versions, we sought to achieve similar rates of licking and reward delivery. Therefore, mice in this version could obtain rewards in a hidden ‘response window’ (a 1500 ms time interval in which either left or right licks could be emitted to acquire reward; same duration as MST, below). This response window was temporally decorrelated from the stimuli^67^. Mice thus licked spontaneously at the two spouts and received occasional rewards. Mice were exposed 2-5 days to this behavioral task before any experiments.

##### *UST: Unisensory version* (N=4/28 animals)

In this version, only visual change was associated with reward availability. Mice were trained to respond to the visual changes only. Continuous auditory stimuli and changes were presented throughout training and recording sessions with the same statistics as the multisensory version, but were not associated with reward and were temporally decorrelated from the task-relevant visual trials. Given that only one side was rewarded in this version, spontaneous licking to this side had a higher probability of being rewarded and therefore the response window was shortened to 1000 ms (i.e., in this window, licks could be produced to acquire reward).

##### *MST: Multisensory version* (N=17/28 animals)

In this version visual and auditory change were both associated with reward availability. Mice were trained to respond in a lateralized manner to each modality: lick to one side to report visual changes, to the other side in case of auditory changes (modality-side pairing was counterbalanced across mice). Therefore, in this version, subjects had to simultaneously monitor both the auditory and visual modality, detect changes in a feature and discriminate the modality in which the change occurred. In other words, mice were required to identify the sensory modality in which a change occurred.

#### Training stages

For each trained modality (vision in UST, vision and audition in MST), training occurred in steps. In the first stage learning was facilitated by (1) only including the easiest trial type (maximally salient trials: 90 degrees orientation change for the visual domain^68^ and 4kHz or ½ octave – in earlier and later cohorts, respectively – for the auditory domain), (2) additional instantaneous changes to increase saliency, (3) a passive reward on the correct side if the animal did not respond within 900 ms, and (4) the opportunity to correct after choosing the incorrect side. These facilitating conditions were phased out throughout the training procedure and trials of varying lower saliency were introduced. Animals were trained until their psychometric curve in the target modalities reached a plateau. For the MST version, animals were first trained in one modality, then the other, after which they were combined (the order of modalities was counterbalanced across mice).

Trials types were pseudorandomly presented (block-shuffled per 10 trials, 10% of trials were catch trials, thus without a stimulus change, 41% visual trials, 41% auditory trials, 8% multimodal trials - see below). The inter-trial interval was taken randomly from an exponential distribution with a mean of six seconds (minimum 3 and maximum 20 seconds). Directly after a stimulus change, a response window of 1500 ms followed in which either left or right licks could be emitted to acquire a reward. Licks during the first 100 ms were not counted as these occurred too early to be considered part of a stimulus-response sequence. The first lick after this ‘grace period’ was registered as the animal’s choice and correct licks were directly rewarded. To counter any bias in MST mice, if the fraction of licks to one spout out of all licks in the last 10 trials was above 90%, the next trial was selected with a 95% probability to be of the other modality. As visual and auditory feature changes were associated with conflicting motor actions (only in the multisensory version of the task), a multimodal trial (simultaneous audiovisual change) would present the animal with conflicting signals. We introduced these conflict trials in a subset of sessions, but these trials were not included in the current analyses.

For each trained animal (before any recordings) we implemented three behavioral sessions in which we presented five levels of auditory and visual saliency that spanned the perceptual range to establish the perceptual sensitivity of each mouse. We fit the concatenated data of these three sessions with a cumulative normal distribution per modality with four free parameters^69^:

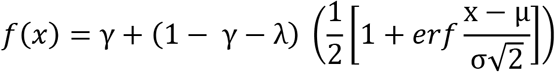

Here, γ describes the false alarm rate (spontaneous licks during catch trials), λ the lapse rate (misses at maximal saliency), μ the mean (perceptual threshold), and σ the standard deviation (sensitivity to variations of stimulus intensity). Having established the psychometric function per mouse, we took four levels of saliency per modality at fixed points along the psychometric function: subthreshold (μ-σ; *sub*), threshold (μ; *thr*), suprathreshold (μ+σ; *sup*), and maximal saliency (*max*). The visual threshold ranged from 4-12 degrees, and the auditory threshold from 10-100 Hz (frequency version) or 1/64 - 1/16 partial octave (octave version) (see Ext. Data Fig. 1). This analysis was purely performed to select stimulus intensities of equal subjective saliency across mice for the experiments. All other analyses were based on fitting the behavioral data with a psychometric signal detection model (see below).

In recording sessions, we limited conditions to sample sufficient trials per modality x feature x saliency x choice combination. First, we only used two levels of change: threshold and maximal saliency. For NE mice and auditory conditions in UST mice, we used threshold values that matched those from trained animals. Second, we only used four orientations or tones. Specifically, this means that stimuli jumped between A, B, C, and D, where distance AB and CD are around threshold and distance AC and BD are maximal. An example stimulus set for a mouse with a visual threshold of 7° and an auditory threshold of 1/32 octave was therefore for the visual domain: A=100°, B=107°, C=190°, D=197°, and for the auditory domain (in Hz): A=2^13.25^, B=2^13.25+1/32^, C=2^13.75^, D=2^13.75+1/32^.

### Behavioral task B: Visuotactile Side Detection

As in paradigm A, mice were trained in on one of two versions of a visuotactile detection task: a multisensory version, where both visual and tactile modalities were informative on the side that needed to be chosen to acquire reward (MST) and a unisensory version, where the tactile modality was present as well, but only the visual modality was informative (UST).

#### Visual Stimuli

Visual stimuli consisted of square-wave drifting gratings, with a temporal frequency of 1.5 Hz, a spatial frequency of 0.025 cycles per degree and 30 degrees orientation. The contrast of the gratings was modulated per trial to control detection difficulty. Visual stimuli were generated in Octave using Psychtoolbox3 and were presented monocularly at >24 degrees from azimuth on each side^70^ with a gamma-corrected 18.5-inch monitor at a frame rate of 60 Hz and a distance to the eye of 18 cm.

#### Tactile Stimuli

Tactile stimuli consisted of a single deflection of the whisker pad using a piezoelectric bender (PL128.10, Physik Instrumente) coupled to a 5 cm long pipette ending on a 5×5 mm patch of Velcro. A voltage driver (E650, Physik Instrumente) and an RC filter were used to produce a backward deflection of the bender with an exponentially decaying speed (τ=72 ms) during 360 ms, followed by a forward deflection with the same characteristics. The amplitude of the deflection was modulated to control detection difficulty. Elicited whisker deflection angles ranged from 0 to 3.6 degrees. For both visual and tactile stimuli, stimulus intensity was adjusted individually to match the desired saliency.

#### Versions of task B

##### UST: Unisensory version

Visual and/or tactile stimuli were presented to either the right or left side of the animal. To obtain a reward, mice had to detect the side where the visual stimulus was presented and lick the spout at the corresponding side. In this version, only the visual modality was informative on reward availability. Tactile stimuli were delivered but not associated with reward and tactile and visual stimulus sides were decorrelated. The inter-trial interval was drawn from an exponential probability distribution with a mean of 4 seconds (minimum 3, maximum 7; with a 22% chance of catch trial (no stimulus, no reward) and a maximum of two catch trials in a row, a mouse could wait up to 21 seconds before another stimulus was displayed). Visual and/or tactile stimuli were presented for 1 second. In a multisensory trial (not analyzed here), the tactile stimulus was presented with a lag of 70 ms after the visual stimulus onset (similar to ^42^). Licks were only rewarded in the interval of 140-1000 ms after stimulus onset. While a correct lick triggered reward delivery, an incorrect lick (i.e., to the wrong side) terminated the trial and aborted stimulus presentation. Trials were pseudo-randomly generated by blocks of 60 with 22% catch trials, 12% tactile-only trials, 53% visual-only trials, and 13% multisensory trials.

##### MST: Multisensory version

In this version, both visual and tactile modalities were informative on reward availability. In multisensory trials, visual and tactile stimuli were presented on the same side. Overall, task B required the mouse to follow an Inclusive-Or rule (lick to the side with either a visual or tactile stimulus, or a compound stimulus in both modalities). During training, mice first learned to detect tactile stimuli. Multisensory trials were then added and finally, visual-only trials were introduced so that mice could eventually detect visual and/or tactile modalities. Since tactile trials were rewarded, to keep the reward/no-reward balance, we increased the number of tactile trials: 25% catch, 25% visual-only, 25% tactile-only, 25% multisensory. Otherwise, both unisensory and multisensory task versions had the same parameters.

### Imaging, Optogenetics, and Electrophysiology

#### Intrinsic Optical Imaging

To localize the primary visual cortex in task B experiments, we performed intrinsic optical imaging (IOI) under lightly anesthetized conditions (0.7-1.2% isoflurane). A vasculature image was acquired under white light before starting the imaging session. During IOI, the cortex was illuminated with monochromatic 630 nm light. Images were acquired using a cooled 50 Hz CCD camera connected with a frame grabber (Imager 3001, Optical Imaging Inc, Germantown, NY, USA), defocused about 500-600 μm below the pial surface. Visual stimulation consisted of square-wave drifting gratings as described in^12^, presented in the right visual hemifield. Image processing was carried out as described in^12^.

#### Viral injection

Mice were subcutaneously injected with the analgesic buprenorphine (0.025 mg/kg) and maintained under isoflurane anesthesia (induction at 3%, maintenance at 1.5–2%) during surgery. We performed small craniotomies (±100 μm) over V1 using an ultrafine dental drill and inserted a glass pipette backfilled with AAV2.1-EF1a-double floxed-hChR2(H134R)-EYFP-WPRE-HGHpA (titer: 7×10^12^ vg/mL, 20298-AAV1 Addgene). In total 50 nL was injected in V1 (bilateral binocular V1 for Task A and unilateral V1 for Task B) at 700 μm and 400 μm below the dura (25 nL per depth) using a Nanoject pressure injection system (Drummond Scientific Company, USA).

#### Optogenetics

In a random subset of trials (50% of trials for task A, 25% for task B) photostimulation started at stimulus onset (early inactivation) or was delayed (late inactivation). For the MST version of task B, early and late inactivation took place in separate sessions. Late inactivation occurred after 200 ms in Task A and 240 ms in Task B. Photostimulation continued until the animal made a choice. We interleaved sessions in which we positioned the fiber over V1 with control sessions in which we either positioned the optic fiber over area S1 (where no virus was injected) or at the head-bar. To locally photostimulate V1, a 473 nm laser (Eksma Optics, DPSS 473nm H300) was connected to one or two fiber-optic cannulas (ID 200 um, NA 0.48, DORIC lenses) that were positioned directly over the thinned skull at the area of interest (bilateral V1 for Task A and unilateral V1 in Task B). Light delivery was controlled by a shutter (Vincent Associates LS6 Uniblitz) with variable pulse and interpulse duration with an average of 20 Hz and 75% duty cycle (Task A) or with 10 ms pulses sequentially interleaved by 20 ms and 30 ms (~72% duty cycle, Task B). The shutter was located in a sound-insulated box distal from the experimental setup. As we simultaneously performed extracellular recordings in V1 of all mice, we adjusted laser power for each animal to the minimum power that maximally inhibited neural activity. This was commonly 2-7 mW/mm^2^ at the cortical surface (2-15 mW15mW at the tip of the fiber, placed 0.5-2 mm above the cortical surface), corresponding to an effective 0.5-1.8 mW/mm^2^ (due to 72-75% duty cycle), which is below the levels that produce unwanted heating in tissue^17,71^.

To prevent light from reaching the eye of the mouse, the cannulae were sealed with black tape, leaving only the tip exposed. Furthermore, sessions with optogenetic manipulation were performed in an environment with ambient blue light. Even though we implemented these measures, we observed an increase in false alarms in some mice in task A. This suggests either that mice could perceive the laser, or that our manipulation evoked perceptual changes that were reported as a trial. We therefore verified (1) that our main effect of late silencing was not explained by a change in criterion (see Behavioral Analysis Task A), (2) positioned the fiber over uninfected somatosensory cortex (S1), and (3) performed the same optogenetic experiments in a second visuotactile paradigm where we did not have an increase in False Alarm responses by photoinactivation of V1.

#### Extracellular recordings

Mice were subcutaneously injected with the analgesic buprenorphine (0.025 mg/kg) and maintained under isoflurane anesthesia (induction at 3%, maintenance at 1.5–2%) during surgery. We performed small (about 200 μm) craniotomies over the areas of interest (up to 6 per animal) using a dental drill. The recording chamber was sealed off with silicon elastomer and the mice were allowed to recover for 24h.

Extracellular recordings were performed on consecutive days with a maximum of 4 days to minimize damage to the cortex. Microelectrode silicon probes (NeuroNexus, Ann Arbor, MI – 4 types of either 32 or 64 channels were used, catalog numbers A1×32-Poly2-10mm-50s-177, A2×16-10mm-100-500-177, A4×8-5mm-100-200-177, A1×64-Poly2-6mm-23s-160) were slowly inserted in the cortex until all recording sites were in contact with the tissue. V1 was approached perpendicularly to the cortical surface. The medial prefrontal cortex, primary auditory cortex, and posterior parietal cortex were also recorded, but data from these areas were not analyzed here. After insertion, the exposed cortex and skull were covered with 1.3-1.5% agarose in artificial CSF (125 mM NaCl, 5 mM KCl, 1.3 mM MgSO_4_, 2.0 mM NaH_2_PO_4_, 2.5 mM CaCl_2_, pH 7.3) to prevent drying and to help maintain mechanical stability. The probe was left in place for at least 15 minutes before recording to allow for tissue stabilization. Electrodes were dipped in DiI (ThermoFisher Scientific) during the final recording session allowing better post hoc visualization of the electrode tract. The ground was connected to the head bar and the reference electrode to the agarose solution. Neurophysiological signals were pre-amplified, bandpass filtered (0.1 Hz to 9 kHz), and acquired continuously at 32 kHz with a Digital Lynx 64/128 channel system (Neuralynx, Bozeman, MT).

Spike sorting of data acquired during task B was done as previously described^42^ and only units having less than 1% of their spikes within a 1.5 ms refractory period were kept. For task A we used Klusta and then manually curated with the Phy GUI^72^. Before spike sorting the median of the raw trace of nearby channels (within 400 μm) was subtracted to remove common artifacts. Each candidate single unit was inspected during manual curation based on its waveform, autocorrelation function, and its firing pattern across channels and time. Only high-quality single units were included, defined as having (1) an isolation distance higher than 10 (cf.^73^) (2) less than 0.1% of their spikes within the refractory period of 1.5 ms^74,75^, (3) spiking present throughout the session. Neurons were deemed stably present if they had spikes in more than 90 out of 100 time bins during the entire session.

#### Recording depth estimation

The estimation of the laminar depth of the electrodes in V1 was based on three aspects. First, we computed the power in the 500-5000 Hz range to localize layer 5 with the highest MUA spiking power^76^. Second, we showed contrast-reversing checkerboards before each recording session and computed the current source density profile to estimate layer 4 with the earliest current sink, as previously described^77^. Lastly, this was aligned with the depth registered when the silicon probes were lowered from the dura. The granular layer was taken to span from 400 to 550 μm from the dura.

#### Video monitoring

In Task A, the left eye (ipsilateral to the hemisphere of recording) was illuminated with an off-axis infrared light source (six infrared LEDs 850 nm) adjusted in intensity and position to yield high contrast illumination of both the eye and whisker pad. A frame-grabber acquired images of 752×582 pixels at 25 frames per second through a near-infrared monochrome camera (CV-A50 IR, JAI) coupled with a zoom lens (Navitar 50 mm F/2.8 2/3” 10MP) that was positioned at approximately 30 centimeters from the mouse.

To extract pupil variables^55,78^ we trained DeepLabCut^79^ on 300 frames from 15 video excerpts of 1-2 minutes with varying pupil size, illumination, contrast, imaging angle, and task conditions. We labeled the pupil center and 6 radially symmetric points on the edge of the pupil. An ellipsoid was fit to these 6 outer points. The pupil center was taken as the center of the ellipsoid and the pupil area as the ellipsoid area from the fitted ellipse parameters. Single poorly fit frames were replaced by the running median (10 frames). We z-scored the total session trace.

#### Histology

At the end of each experiment, mice were overdosed with pentobarbital and perfused (4% paraformaldehyde in phosphate-buffered saline), and their brains were recovered for histology to verify viral expression and placement of silicon probes in V1. We cut coronal 50 μm sections with a vibratome, stained them with DAPI, and imaged the mounted sections. For flattened cortical sections (e.g. Fig. 3b) we first removed subcortical tissue and flattened the cortical sheet of each hemisphere between glass slides by applying pressure overnight before sectioning 100 μm slices with the vibratome (as described previously^80^). For coronal sections, area borders were drawn by aligning and overlaying the reference section from the atlas^81^. For flattened cortical sections, area borders were drawn based on cell densities aligned to reference maps^82^.

### Data analysis

Unless otherwise stated, all data were analyzed using custom-made software written in MATLAB (The MathWorks, Natick, MA).

#### Behavioral analysis - Task A

Sessions were terminated when the animal did not respond for 20 trials and these last 20 trials were discarded from analyses. Sessions in which the hit rate for maximal auditory and visual changes was below 30% were excluded.

Behavioral response rates in task A were fit with a multi-alternative signal detection model^35^. This model extends signal detection theory^40^ and aims to accurately and parsimoniously account for observer behavior in a detection task with multiple signals. In this model, the decision is based on a bivariate decision variable whose components encode sensory evidence in each modality. Decision space is partitioned into three regions (no response: neither evidence is strong enough; auditory response, and visual response). In a given trial, the observer chooses to report visual or auditory stimuli if the decision variable exceeds a particular cutoff value, the ‘‘criterion’’ for each signal (the animal’s internal signal threshold for responding, in terms of signal detection framework). We fit two versions of this model. In sessions with two levels of saliency (threshold and maximum), we fit the d-prime (d’) and criterion (c) to the behavioral response rates separately for each stimulus change intensity. This consists of fitting four free parameters (d’ and c for each modality). In sessions with four or five levels of saliency per modality, we fit the behavioral response rates by fitting a criterion per modality and a d-prime for each saliency, which is described by a psychophysical function (three-parameter hyperbolic function). The d-prime at each saliency level follows from:

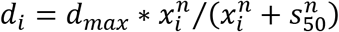

where *d_max_* is the asymptotic d-prime, *s_50_* is the stimulus strength at 50% of the asymptotic value, *n* is the slope of the psychometric function and *x_i_* is the amount of change. This consisted of fitting a total of 8 free parameters: *d_max_*, *n*, *s_50_*, and *c* for each modality. We refer the reader to Sridharan *et al.* (2014) for a detailed description of how the d-prime and criterion subsequently relate to response rates^35^. Single session fits where visual threshold was below 1 degree or above 45 degrees were excluded (average threshold ±6 degrees, n=3/179 sessions excluded).

Catch trials during the tasks served to measure baseline lick responses. As there were no stimulus changes during the inter-trial interval (visual and auditory stimuli continued to be presented throughout the session, similar to catch trials), we used long inter-trial intervals to insert additional artificial catch trials during offline analysis to achieve increased balance across trial type conditions. We controlled for temporal expectation and inserted additional catch trials only at time points conforming to the original inter-trial interval statistics.

After analyzing the effects of early and late V1 silencing on audiovisual change detection on the full dataset, we focused in a subsequent analysis on the relationship between reaction time and the effect of late silencing (Fig. 3i,j, Ext. Data Fig. 6d,e,f). Here, we focused on sessions in which V1 early silencing was effective (minimum 50% reduction in d-prime on maximal visual change, 59/81 sessions; results were robust to variations of this criterion, 25% reduction, r=0.199, p=0.049; 75% reduction, r=0.511, p=0.001). This threshold was implemented to test if late silencing was effective specifically within those sessions in which the optogenetic manipulation demonstrably impaired visual detection (thus exploiting an internal control).

#### Behavioral analysis - Task B

Behavioral data in task B was fit with a multinomial logistic regression, as described in^83^. The probabilities of right choice (*p_right_*), left choice (*p_left_*) and no choice (*p_no-go_*) were set by:

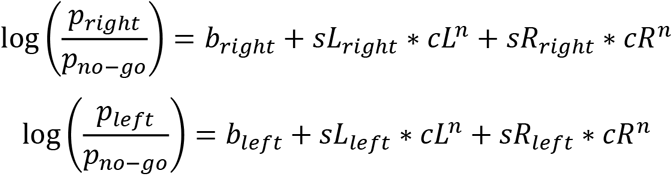

Where *b* is a bias parameter, *sL* and *sR* are the sensitivity to stimulus evidence to the left and right side respectively, *cL* and *cR* are the stimulus intensity to the left and right side respectively (contrast for vision, deflection angle for somatosensation), *n* is an exponent parameter between ranging between 0 and 1 to allow for saturation.

The model was fit to individual mice, with all sessions pooled together. However, per mouse, visual behavior (visual-only trials) and tactile behavior (tactile-only trials) were fit separately. The model was fit using Matlab’s *mnrfit* and maximum likelihood estimation. To quantify behavioral performance, we computed d-prime (d’) as:

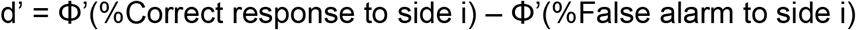

Where Φ’ is the normal inverse cumulative distribution function.

#### Electrophysiological data processing

To visualize the effect of photostimulation on spiking activity at a single electrode channel the raw signal was high-pass filtered (500 Hz, 4^th^ order Butterworth filter). To compute firing rates, spikes (following spike detection and sorting) were binned in 10 ms bins and convolved with a causal Half-Gaussian window with 50 ms standard deviation, unless stated otherwise. Wherever firing rate was z-scored, the mean was subtracted and divided by the standard deviation of the baseline period (−1 to −0.2 seconds before stimulus). For Figure 3d the firing rate was only normalized to the baseline to quantify the relative reduction in firing rate by optogenetic inhibition. For Figure 5g,h the standard deviation of the convolutional window was reduced to 10 ms to enhance temporal resolution. For computing noise correlations the standard deviation of the convolutional window was increased to 100 ms to increase noise correlation estimates. For Figure 4c and Ext. Data Fig. 9c-d, neurons with an average z-scored firing rate that exceeded 2 standard deviations at any point during the stimulus epoch of any visual trial condition were considered responsive and included. Neurons in sessions lacking any of the compared conditions were excluded.

#### Encoding model of single neuron firing rates

To quantify single neuron encoding of different task variables, we constructed a kernel-based Poisson regression model. This encoding model allowed us to model, for single neurons, the time-dependent effects of all measured variables related to the task and the animal’s behavior simultaneously on single-trial neuronal activity^37,38^. This approach is particularly useful to disentangle the unique contribution of experimenter-controlled task events and self-timed behavioral events to variability in firing rates across the neuronal population.

##### Construction

For each neuron, we constructed a design matrix based on five sets of variables; visual, auditory, hit/miss, movement, and arousal variables. Binary variables (all except pupil size) were modeled with a series of temporal basis functions (raised cosines) that spanned the relevant epoch of influence. The number and temporal distribution of these basis functions were selected to maximize the cross-validated explained variance (see below). For the sensory predictors, we used two kernels with 100 ms standard deviation that spanned the first 200 ms post-stimulus to capture the early spiking activity and 10 kernels with 200 ms standard deviation that spanned from 0 to 2000 ms post-stimulus to capture the late, sustained response. We found that making a separate predictor set per combination of orientation x amount of change produced the highest quality fit as it simultaneously took into account the selectivity of neurons for orientation and saliency. This therefore resulted in (2 + 10 basis functions) x 2 (modalities) x 2 (levels of change) x 2 (grouped post-change features) = 96 predictors. For hit/miss variables we used 10 temporal basis functions with 200 ms standard deviation that spanned from 0 to 2000 ms relative to stimulus change in hit trials (visual hit, audio hit) and 10 predictors that spanned −500 ms to +1500 ms relative to reward (20 predictors for hit/miss). For movement variables, we used three basis functions that spanned −200 to +400ms relative to each lick, split by side (6 predictors). To capture arousal effects, the z-scored pupil area was included in the predictor set: with original timing and two temporal offsets (−800 ms and −400 ms) to account for the delayed relationship of brain state to pupil size (e.g.^84^; this equals 3 predictors). We included one whole-trial variable that scaled with the within-session trial number. This full model summed up to 126 predictors. We compared the performance of this model to a null model, with one predictor (a random variable). For convenience, all predictors were normalized to their maximum values before being fed into the model.

##### Fitting

We fitted the encoding model to each neuron’s activity individually, using the *glmnet* package in Matlab^85^ with elastic-net regularization and a Poisson link function, which involves setting three hyperparameters. First, we chose elastic net mixing parameter α = 0.95 to allow for a small number of uncorrelated informative predictors to be favored. Second, model performance was trained and tested on separate data with 5-fold cross-validation. Third, to maximally punish weights without losing model fit quality, regularization parameter lambda was maximized while keeping the cross-validated error within one standard error of the minimum (lambda_1se in *glmnet*). Because very sparsely firing neurons produced fitting difficulties, only neurons with a session-average firing rate >0.5 Hz were included.

##### Evaluation

We quantified the model performance by assessing the 5-fold cross-validated Explained Variance (EV) by the predicted firing rate based on the random or full model, or a subset of predictors from the full model. Explained Variance was calculated as:

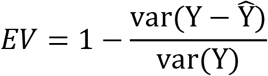

where Y is the original firing rate and Ŷ the estimated firing rate. Explained Variance was computed in two ways. First, we computed EV over all concatenated firing rate bins (over all single trials; −0.5 to 2.5 seconds relative to stimulus change). Second, we computed EV on the concatenated firing rate bins of the average firing rate for five trial-type x choice conditions that captured most trial counts^20,38^ (85% of all trials). To compute EV over time we computed the explained variance over all concatenated time bins at a specific moment relative to stimulus onset.

#### Decoding single neuron activity

To identify which variables were encoded in single neurons we used ROC analysis^40^ and identified how well an external observer could discriminate variables from the firing rate at single time points. We computed the area under the ROC curve (AUC) for the firing rate distributions between two selections of trials. Each class had to have at least 10 trials. AUC values are in the range of 0 to 1 and were rectified to their corresponding values in the range between 0.5 and 1. We investigated three types of coding and for each of these we analyzed threshold change and maximum change trials separately:

*Visual Orientation*: We grouped the pairs of post-change orientations that were close to each other (e.g. A and B oriented at 90 and 97 degrees, see above) and thus compared the firing rate distributions of A&B versus C&D for threshold and maximal change trials separately.
*Occurrence of visual change*: We tested whether single neuron firing rates discriminated between visual and catch trials.
*Hit/miss*: To identify significant coding of the detection of a visual stimulus we compared firing rate distributions within visual trials for hits and misses.

To determine the significance of AUC values at each time bin and for each comparison, we performed a permutation test by shuffling the class labels across trials 1,000 times. If the unshuffled AUC value exceeded 99% of the shuffled distribution (P < 0.01) this was deemed significant. This yielded an AUC value for each neuron for each time bin for each type of coding and each of these values its significance by the permutation test.

To compare coding dynamics across cohorts, we normalized the fraction of significantly coding neurons by subtracting the baseline fraction (average over −0.5 to 0 seconds) and dividing by the maximum. Each condition was only normalized to maximum if the fraction of significantly coding neurons increased at least 10% over baseline.

To determine the onset of significant coding we tested when the fraction of coding neurons increased significantly above a multiple of standard deviations of the coding fraction during baseline. We report results at a threshold of 2 standard deviations (Zscore > 2), but the results were robust to variations in threshold (e.g. 1 or 3 standard deviations). To estimate the reliability of the onset of coding and the relationship of hit/miss coding to reaction time, we bootstrapped by resampling from the total neuronal population (n=1000 bootstraps). To investigate the relationship between the onset of hit/miss coding and reaction time we used a linear regression, which revealed a systematic relationship between the timing of hit/miss coding and reaction time (Fig. 2k). To estimate by how much hit/miss coding preceded reaction time we used two measures. First, we fixed the slope of the regression fit at 1 and found an offset of 278 ms. This was similar across variations of threshold (Z > 1: 288 ms, Z > 3: 250 ms). Second, for each bootstrap, we computed the onset of hit/miss-coding according to the fit parameters for the average reaction time, which was on average 266 ms before the reaction time.

For laminar depth localization of coding dynamics, neurons were binned according to their recorded depth in 50 μm bins spanning from 0 to 1150 μm below the dura. The fraction of neurons coding for each variable at this depth was computed for each time point (25 ms temporal bins). This heatmap was convolved for display purposes with a two-dimensional Gaussian (standard deviation of 1.3 bins – temporal and spatial). For statistical comparison across laminar zones, the fraction of coding neurons was computed for each session (if at least 10 neurons were recorded at this depth to estimate coding fraction reliably) in supragranular, granular, or infragranular layers (granular layer: 400-550 μm from dura). As sensory and hit/miss-coding was present in different temporal epochs these were included for statistical comparison (Orientation 0-1000 ms, Visual occurrence: 0-200 ms, Hit/miss: 200-1000 ms, relative to stimulus change).

#### Population coding analysis

To decode visual stimulus orientation, we departed from the four orientations and grouped the two pairs of orientations close to each other to obtain a two-class classification problem (AB vs CD, see above). Decoding was performed on recordings that contained at least 15 neurons and in which at least 20 trials per orientation pair were available. We equalized the number of neurons across sessions by randomly drawing 10 neurons from all sessions with more than 10 units. Spikes were binned using a sliding window of 200 ms with 50 ms increments, excluding time bins that contained both pre and post-stimulus spikes. Decoding was performed using a random forest classifier with 200 trees, as implemented in Scikit-learn^86^, and we employed a 5×5 cross-validation routine with stratified folds (cf. Bos et al. 2020). The average accuracy obtained in the cross-validation routine was corrected by subtracting the average accuracy on 50 surrogate datasets in which the orientation labels were permuted across trials to obtain the improvement in decoding accuracy beyond chance level.

#### Noise Correlations

To investigate correlated activity across the population we computed pairwise correlations on the binned spike counts (10 ms bins, time range: −1000 to +1500 ms relative to stimulus change) after subtracting the average stimulus-driven response. First, for each neuron the trial-mean firing rate over time was subtracted for all subsets of trials of interest (per orientation). Next, the Pearson’s correlation coefficient was computed between the residual rates for each simultaneously recorded neuronal pair for each time bin. Neuronal pairs for a given condition were included if they were sampled in more than 10 trials. We also computed pairwise correlations aligned to lick onset and thus subtracted mean activity related to lick-related modulation of firing rate. We note, however, that the term ‘noise correlations’ is conventionally reserved for the correlations on residual rates after subtracting the mean stimulus-evoked activity (rather than movement-evoked activity). Note also that drop in noise correlations was not a direct result of overall increased firing rates as we observed no such reduction of noise correlations during early sensory-evoked activity (0 – 200 ms). Neurons were only included if their session average firing rate was above 1 Hz.

#### Statistics

All relevant statistical analyses, *p* values, and *n* sizes are reported in Supplementary Table 1. Data were analyzed using non-parametric tests as a default unless mentioned otherwise. Results with a *p*-value lower than 0.05 were considered significant.

