## Extended Data for "Task complexity temporally extends the causal requirement for visual cortex in perception"

#### Extended Table 1: Statistical tests:

| Fig | Goal | Test | Sample sizes | p-value (significant in bold) |
|---|---|---|---|---|
| 1g | Visual d-prime UST vs MST | Wilcoxon rank sum test with Bonferroni correction for multiple comparisons | 12 UST sessions, 139 MST sessions | p=0.3 |
| 1g | Auditory vs visual d-prime in MST | Wilcoxon signed rank test with Bonferroni correction for multiple comparisons | 139 MST sessions | **p=1.86e-6** |
| 1h | Visual threshold UST vs MST | Wilcoxon rank sum test | 12 UST sessions, 139 MST sessions | p=0.86 |
| Not shown | Visual sensitivity UST vs MST | Wilcoxon rank sum test | 12 UST sessions, 139 MST sessions | **p=0.07** |
| 1i | Linear dependence between visual saliency and RT (UST) | Pearson correlation | 484 trials | **r=-0.123,p=0.00667** |
| | Linear dependence between visual saliency and RT (MST) | | 3381 trials | **r=-0.190, p=6.4e-29** |
| 1i | Auditory vs visual RT (MST) | Wilcoxon rank-sum test | 3381 visual trials, 4229 auditory trials | **p=3.05e-230** |
| 1i | Visual RT for UST vs MST for each saliency level | Wilcoxon rank-sum test with Bonferroni correction for multiple comparisons | Subthreshold:<br>71 UST trials, 483 MST trials | **p=2.46e-04** |
| | | | Threshold:<br>115 UST trials, 748 MST trials | p=0.83 |
| | | | Suprathreshold:<br>130 UST trials, 933 MST trials | **p=1.46e-05** |
| | | | Max:<br>168 UST trials, 1217 MST trials | **p=3.00e-03** |
| 2i | Fraction of neurons coding (significant AUC) different across layers | Two-sided Wilcoxon rank-sum test between each pair of layers with Bonferroni correction for multiple comparisons<br><br>(Only significant reported, rest p>0.05) | Visual orientation; no significant layer differences. | p>0.05 |
| | | | Visual change occurrence; 4 sessions with enough supragranular neurons, 14 sessions with enough infragranular neurons | **p=0.018** |
| | | | Hit/miss threshold; 3 sessions with enough granular neurons, 15 sessions with enough infragranular neurons | **p=0.039** |
| | | | Hit/miss maximal; 3 sessions with enough granular neurons, 15 sessions with enough infragranular neurons | **p=0.027** |
| 2j | Earliest time point of increase in coding fraction | Fraction significant neurons exceeds 2 std above baseline (-1000 to 0 ms). This corresponds to a one-sided t-test with p < .02275. Very similar results were obtained with a threshold of 1 or 3 std above baseline. | NE: 116 neurons<br>UST: 128 neurons<br>MST: 306 neurons | **Z>2** |
| 2j | For each variable, the difference in latency to coding between cohorts | Bootstrap test (n=1000 resamples). If the difference between bootstrap distribution exceeded the 97.5 percentile this was deemed significant. This corresponds to a two-sided p-value of 0.05. | | **Hit/miss coding latency threshold changes between UST and MST p<0.05.** Rest p>0.05. |

| | | | | |
|---|---|---|---|---|
| 2k | Linear dependence earliest increase in hit/miss coding and RT | Pearson correlation | 4 conditions (2 saliencies, UST and MST) | **r=0.989, p=0.01** |
| | | | Bootstrap results: (Mean and 95% CI) | Slope: 1.58 (0.27-2.52)<br>Offset: -569 ms (-985 to -10). |
| 3g-h | Effect of inactivation on discrimination performance (d-prime) comparing early or late inactivation versus control trials for different saliencies, modalities, and cohorts | Wilcoxon signed-rank test with Bonferroni correction for multiple comparisons | threshold visual change, UST, Ctrl vs Early<br>18 sessions | **p=0.005540** |
| | | | threshold visual change, UST, Ctrl vs Late<br>18 sessions | p=1.000000 |
| | | | threshold visual change, MST, Ctrl vs Early<br>34 sessions | **p=0.000523** |
| | | | threshold visual change, MST, Ctrl vs Late<br>34 sessions | **p=0.030450** |
| | | | maximum visual change, UST, Ctrl vs Early<br>18 sessions | **p=0.001159** |
| | | | maximum visual change, UST, Ctrl vs Late<br>18 sessions | p=0.536430 |
| | | | maximum visual change, MST, Ctrl vs Early<br>34 sessions | **p=0.000684** |
| | | | maximum visual change, MST, Ctrl vs Late<br>34 sessions | **p=0.006089** |
| | | | auditory change, MST, all comparisons<br>34 sessions | All p>0.1 |
| 3i | Linear dependence median RT and percentage reduction d-prime | Linear regression | Early silencing: 40 conditions (21 Thr and 19 Max) | R=0.048,<br>P=0.865 |
| 3j | Linear dependence median RT and percentage reduction d-prime | Linear regression | Late silencing: 46 conditions (23 Thr and 23 Max) | **R=0.423,<br>P=0.03** |
| 4d | Effect of inactivation on D-prime. Comparing Early or Late inactivation versus control trials for different saliencies, modalities, sides, cohorts. | Wilcoxon signed-rank test with Bonferroni-Holm correction for multiple comparisons | Visual contralateral threshold detection, UST, Ctrl vs Early<br>7 sessions | **P=0.031** |
| | | | Visual contralateral threshold detection, UST, Ctrl vs Late<br>7 sessions | P=0.078 |
| | | | Visual contralateral threshold detection, MST, Ctrl vs Early<br>6 sessions | **P=0.031** |
| | | | Visual contralateral threshold detection, MST, Ctrl vs Late<br>7 sessions | **P=0.008** |
| | | | Visual contralateral maximum detection, UST, Ctrl vs Early<br>7 sessions | P=0.578 |
| | | | Visual contralateral maximum detection, UST, Ctrl vs Late<br>7 sessions | P=0.4375 |
| | | | Visual contralateral maximum detection, MST, Ctrl vs Early<br>4 sessions | P=0.625 |
| | | | Visual contralateral maximum detection, MST, | P=0.094 |

| | | | | |
|---|---|---|---|---|
| | | | Ctrl vs Late<br>7 sessions | |
| | | | Visual ipsilateral threshold detection, UST, Ctrl vs Early<br>7 sessions | P=1 |
| | | | Visual ipsilateral threshold detection, UST, Ctrl vs Late<br>7 sessions | P=1 |
| | | | Visual ipsilateral threshold detection, MST, Ctrl vs Early<br>2 sessions | P=1 |
| | | | Visual ipsilateral threshold detection, MST, Ctrl vs Late<br>7 sessions | P=1 |
| 4e | Linear dependence between percentage reduction d-prime and reaction time | Linear regression | 30 conditions (7 UST thr, 7 UST high, 9 MST low, 7 MST high) | P=0.0274<br>r-squared=0.162 |
| 5b | Pre-stim (-500 to 0 ms) vs post-stim (0 to +500 ms) decoding improvement over chance | Wilcoxon signed-rank test | 11 sessions combined across cohorts | p=0.00195 |
| 5c | Significant decrease in noise correlations versus baseline for visual trials split by choice | Two-sided Wilcoxon rank-sum test against zero with Bonferroni correction for multiple comparisons (n=6) | NE – Miss trials, 2904 pairs | P<1e-4 |
| | | | NE – Hit trials, 1476 pairs | p=1.000 |
| | | | UST – Miss trials, 1930 pairs | p=1.000 |
| | | | UST – Hit trials, 1930 pairs | P<1e-17 |
| | | | MST – Miss trials, 10048 pairs | p=0.332 |
| | | | MST – Hit trials, 9626 pairs | P<1e-18 |
| 5d | Difference in visual hit reaction time between cohorts | Two-sided Wilcoxon rank-sum test | 1821 UST visual hits vs 1691 MST visual hits | p=1.856e-68 |
| 5e | Earliest time point of decorrelation | Earliest time point that noise correlations drop below baseline minus two standard deviations values. This corresponds to a one-sided t-test with p<0.05. Similar results were obtained with more or less strict thresholds. | From 59 sessions:<br>UST: 4730 neuron pairs MST: 17826 neuron pairs | Z<-2 |
| 5f | Linear dependence reaction time and earliest time point of decorrelation | Pearson correlation | 6 condition averages (fast, mid and slow tertiles for UST and MST each) | r=0.960,<br>p=0.002 |
| 5g | Difference in z-scored activity between hit-miss during 100-200 ms window | Two-sided Wilcoxon signed-rank test with Bonferroni correction for multiple comparisons | UST - Thr - 78 neurons | P=0.001 |
| | | | UST - Max - 78 neurons | P=0.029 |
| 5h | Difference in z-scored activity between hit-miss during 100-200 ms window | Two-sided Wilcoxon signed-rank test with Bonferroni correction for multiple comparisons | MST - Thr - 134 neurons | P=0.001 |
| | | | MST - Max - 120 neurons | P=0.34 |
| 5i | Difference in noise correlation for each time bin | Two-sided bootstrapped confidence interval test | 230 UST and MST neurons, 1564 neuron pairs, 1000 bootstraps | Black lines in Figure 5i, indicate P<0.05 |
| Extended Data figures: | | | | |
| S1f | Linear dependence between auditory d-prime and RT | Pearson correlation | 311 conditions (from 139 MST sessions) | R=-0.26,<br>p=3.2997e-6 |

| | | | | |
|---|---|---|---|---|
| S1g | Linear dependence between visual d-prime and RT | Pearson correlation | 356 conditions (from 151 UST and MST sessions) | **r=-0.32, p=3.5410e-10** |
| S2d | Difference in trial-averaged z-scored firing rate in 0 to 200ms post-stimulus window between threshold and maximal visual change conditions per cohort | Two-sided Wilcoxon signed-rank test | 163 NE neurons | **P=3.38e-03** |
| | | | 128 UST neurons | **P=8.42e-06** |
| | | | 525 MST neurons | **P=4.91e-12** |
| S3g | Difference in trial-averaged z-scored firing rate in -300 to 300ms window between lick and no-lick conditions | Kruskal-Wallis test (nonparametric one-way analysis of variance), with Bonferroni posthoc | 163 NE neurons,<br>128 UST neurons<br>525 MST neurons | **p-value in figure, *p<0.05, **p<0.01, ***p<0.001** |
| S3h | Difference in trial-averaged z-scored firing rate in -300 to 300ms window between hit and error conditions for trained UST and MST conditions | Two-sided Wilcoxon signed-rank test with Bonferroni correction for multiple comparisons | Visual Errors vs Hits, n = 128 UST neurons | **P=2.50e-05** |
| | | | Visual Errors vs Hits, n = 525 MST neurons | **P=1.20e-22** |
| | | | Auditory Errors vs Hits, n = 525 MST neurons | **P=8.03e-13** |
| S4b | Difference in explained variance over *single* trials between cohorts over all trials using all predictors | Kruskal-Wallis test (nonparametric one-way analysis of variance), with Bonferroni posthoc | NE vs UST<br>n=116 NE neurons<br>n=128 UST neurons<br>n=272 MST neurons<br>(only neurons >0.5 Hz) | p=1 |
| | | | NE vs MST | **p=0.007** |
| | | | UST vs MST | **p=0.033** |
| S4c | Difference in explained variance over *averaged trial types* between cohorts over all trials using all predictors | Kruskal-Wallis test (nonparametric one-way analysis of variance), with Bonferroni posthoc | NE vs UST<br>n=116 NE neurons<br>n=128 UST neurons<br>n=272 MST neurons | p=0.638 |
| | | | NE vs MST | **p=0.001** |
| | | | UST vs MST | p=0.104 |
| S4e | Explained variance averaged over 0-200ms (early activity) between predictor sets within cohort | Kruskal-Wallis test (nonparametric one-way analysis of variance), with Bonferroni posthoc | n=116 NE neurons<br>n=128 UST neurons<br>n=272 MST neurons | **p-value in figure, *p<0.05, **p<0.01, ***p<0.001** |
| S4f | Explained variance averaged over 200-1000ms (late activity) between predictor sets within cohort | Kruskal-Wallis test (nonparametric one-way analysis of variance), with Bonferroni posthoc | n=116 NE neurons<br>n=128 UST neurons<br>n=272 MST neurons | **p-value in figure, *p<0.05, **p<0.01, ***p<0.001** |
| S6b | Effect of inactivation on discrimination performance (d-prime) comparing early or late inactivation versus control trials for auditory saliencies | Wilcoxon signed-rank test with Bonferroni correction for multiple comparisons | auditory change, MST, all comparisons<br>34 sessions | All p>0.1 |
| S6f | Linear dependence median RT and percentage reduction in criterion | Linear regression | Late silencing: 87 conditions (45 Thr and 42 Max) | R=-0.25, P=0.291 |
| S6g | Late silencing delays reaction times: difference in reaction time between control and late silencing visual hits | Wilcoxon rank-sum test | *maximal* changes:<br>Control hits (n=390 trials) and late silencing hits (n=161 trials) | **P=0.024** |
| | | | *threshold* changes:<br>Control hits (n=274 trials) and late silencing hits (n=122 trials) | P=0.177 |
| S7b | D-prime visual and audio change, UST and MST, Ctrl vs Early and Ctrl vs Late | Wilcoxon signed-rank test with Bonferroni correction for multiple comparisons (n=3) | 13 UST sessions, 16 MST sessions | Non-significant, all comparisons p>0.05 |

| | | | | |
|---|---|---|---|---|
| S8a | D- prime maximum, visual UST vs MST | Wilcoxon rank sum test | UST: 4 mice x 7 sessions x 2 sides<br>MST: 4 mice x 15 sessions x 2 sides | P=0.147 |
| **S8b** | Visual detection threshold, UST vs MST | Wilcoxon rank sum test | MST: 4 mice x 2 sides<br>UST: 4 mice x 2 sides | p=0.701 |
| **S8c** | UST Visual RT, Thr vs Max | Wilcoxon rank sum test | 14 values (7 sessions, left and right RTs) | **p=0.0159** |
| | MST Visual RT, Thr vs Max | | 30 values each (15 sessions, left and right) | **p=0.0099** |
| | MST Tactile RT, Thr vs Max | | 30 values each (15 sessions, left and right) | p=0.3751 |
| | Visual RT (max and thr), UST vs MST | | 28vs60 values | **p=0.011** |
| **S8d** | D-prime tactile contralateral, threshold, MST, Ctrl vs Early | Wilcoxon signed-rank test with Bonferroni-Holm correction for multiple comparisons | 6 sessions | p=0.264 |
| | D-prime tactile contralateral, threshold, MST, Ctrl vs Late | | 9 sessions | p=0.667 |
| S8e | UST Visual Contra Thr, Early vs Ctrl | Wilcoxon rank sum test with Bonferroni-Holm correction | 7 sessions | **P = 0.0313** |
| | UST Visual Contra Thr, Late vs Ctrl | | 7 sessions | **P=0.0156** |
| | MST Visual Contra Thr, Late vs Ctrl | | 7 sessions | **P=0.0469** |
| | MST Visual Contra Thr, Early vs Ctrl | | 6 sessions | **P=0.0469** |
| S10a | Significant decrease in noise correlations versus baseline for *audio* trials split by choice | Two-sided Wilcoxon rank-sum test against zero with Bonferroni correction for multiple comparisons (n=6) | NE – Miss trials, 2904 pairs | P=0.59 |
| | | | NE – Hit trials, 2106 pairs | p=0.54 |
| | | | MST – Miss trials, 9310 pairs | p=1.8e-9 |
| | | | MST – Hit trials, 10116 pairs | P=7.3e-16 |
| S10b | Linear dependence reaction time and earliest time point of decorrelation relative to first lick | Pearson correlation | 6 condition averages (fast, mid and slow tertiles for UST and MST each) | r=0.738, p=0.094 |

### EXTENDED DATA FIGURES:

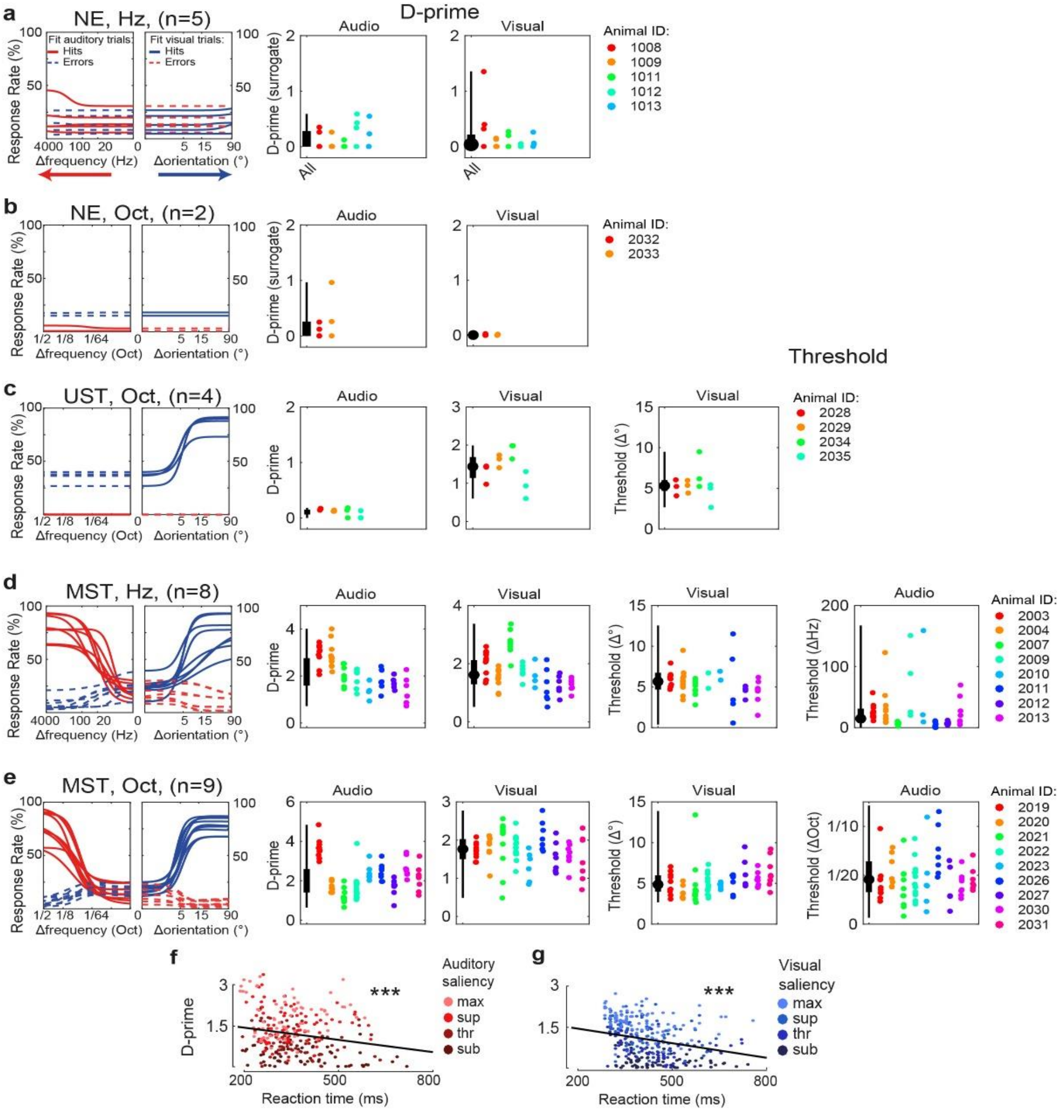


**Extended Data Figure 1: Detailed psychophysical performance across versions of task A.** This figure shows the data for individual animals and individual sessions for each variant of task A. We implemented two versions of auditory stimuli (frequency changes expressed in the amount of Hertz, or octaves, see Methods) and split figures here based on version. The figure follows the same conventions as Figure 1d-f in the main text. Dots in D-prime graphs indicate individual sessions from a single mouse (identified by color). a) Animals (N=5) were used for the noncontingently exposed (NE) variant with auditory changes in Hz. The two leftmost panels show fitted psychometric functions (two-alternative detection model) displaying behavioral response rates at increasing levels of auditory change (left panel) and increasing levels of visual orientation change (right panel). Solid lines are hits and dotted lines are errors. Blue indicates responses to the visual lick spout and red responses to the auditory lick spout. These baseline lick rates after stimulus changes in NE mice result from the animal spontaneously licking (as some licks were rewarded, but not temporally related to the stimuli, see Methods), of which some happen to fall after stimuli ('surrogate hits'). The subpanels to the right side show for each animal (different rainbow colors) and for each session (single dots) the parameters of the single session fits for the asymptotic d-prime for auditory detection (left) and visual detection (right). Within each subpanel, the median and interquartile ranges across all sessions are shown as a boxplot. b-e) Same as **a**, but for the other reward contingencies for task A (UST and MST). For animals trained to report either visual or auditory changes, the detection thresholds are also shown. These include the visual threshold in UST and the visual and auditory threshold in MST. Thresholds for non-rewarded conditions were higher than the maximum saliency or infinite. No mice were trained for the UST variant of the task with auditory changes in Hz. f) In MST animals, D-prime decreases as a function of reaction time both for auditory (upper panel, r=-0.26, p<1e-6). Each dot is one saliency condition within a single session. g) Same as **f**, but for the negative correlation between reaction time and d-prime in all visual conditions across combined UST and MST sessions (r=-0.32, p<1e-10).

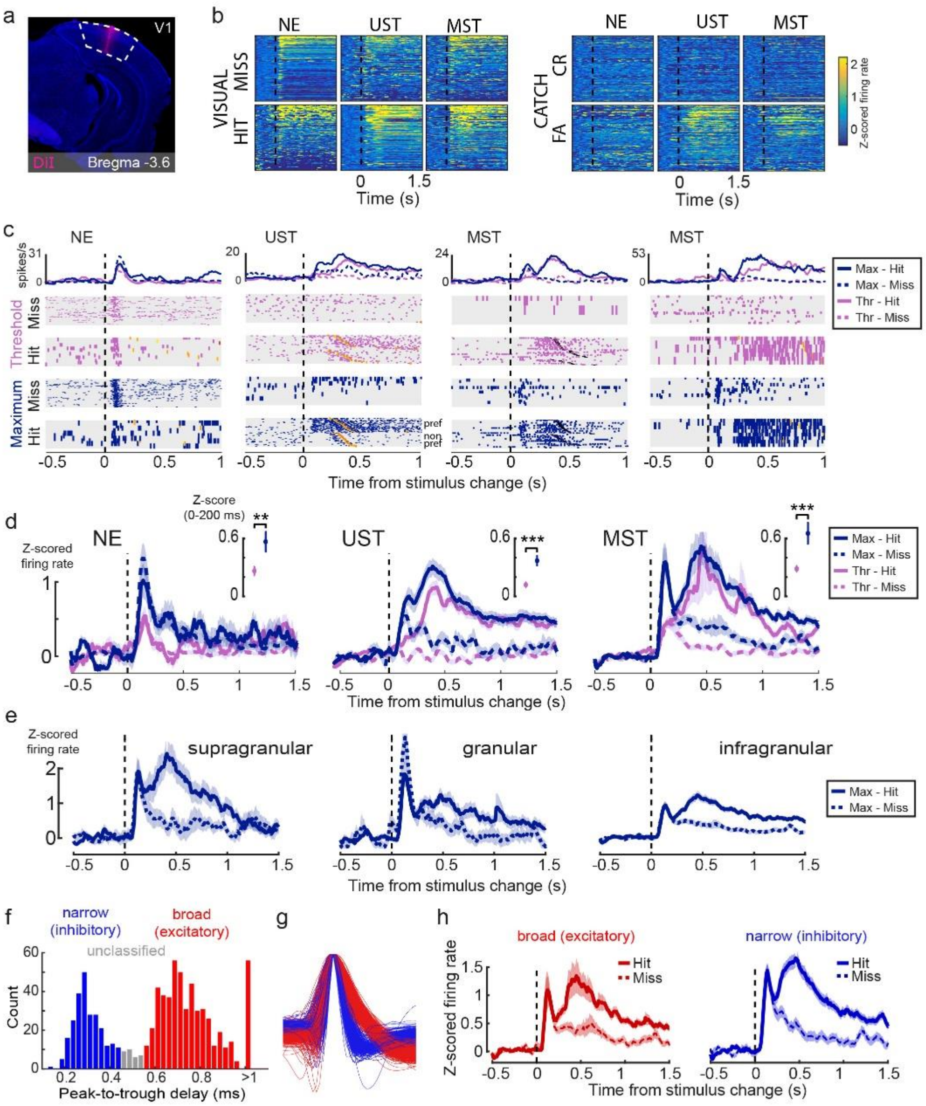


**Extended Data Figure 2: Early and late wave dynamics are present across levels of change saliency and neuron types (Task A).** a) Coronal histological section showing silicon probe tract after recording in V1. b) Heatmaps of trial-averaged z-scored activity of all neurons for the three cohorts for the same conditions as in Figure 2a-c, comparing visual and catch trials split by hit/miss. Note that hits and misses in NE mice are surrogate conditions and are defined post-hoc. Neurons are sorted by their average post-change activity for visual hits (0-500 ms). NE neurons: n=159; UST neurons: n=128; MST neurons: n=510. c) Raster plots for four example neurons showing early sensory-driven transients and late activity both for threshold and maximal orientation changes. Orange ticks indicate first lick and immediate reward delivery. Same conventions as in Figure 2. d) Averaging Z-scored firing rate across all neurons again reveals early sensory-driven activity in all cohorts, while hits are associated with a strong late increase in activity only in UST and MST mice for both visual saliency levels. The amplitude of early sensory-driven activity (average activity 0-200 ms, hits and misses combined) was smaller for threshold than for maximal changes for all three cohorts (shown in insets, NE: $p=3.38*10^{-3}$, UST: $p=8.4*10^{-6}$, MST: $p=4.9*10^{-12}$). Lines and shading indicate mean ± s.e.m. across neurons. e) Same as d, but for UST and MST neurons based on the laminar zone in maximal changes only. Note how early sensory-induced activity is most prominent in the granular and supragranular layers, while late hit/miss modulation is most strikingly different in infra- and supragranular layers. f) We classified single units based on the delay between peak voltage deflection and subsequent trough. The histogram of peak-to-trough delay showed a bimodal distribution (n=816 neurons, all cohorts) allowing clear classification into cell types: narrow-spiking (peak-to-trough delay<0.45 ms; putative inhibitory; blue) and broad-spiking (peak-to-trough delay>0.55 ms; putative excitatory; red). The peak-to-trough delay was capped at 1 ms for neurons whose trough extended beyond the sampled window. Single units with intermediate peak-to-trough values were unclassified. g) Normalized average waveform for all V1 neurons colored by cell type class. h) The dynamics of early and late components were present in both cell classes. This suggests a balanced increase in both excitatory and inhibitory activity upon hits. Z-scored activity averaged over all broad-spiking V1 neurons (left, n=421 neurons) or narrow-spiking neurons (right, n=202 neurons) across UST and MST mice, split by hit/miss response for maximum visual change trials only.

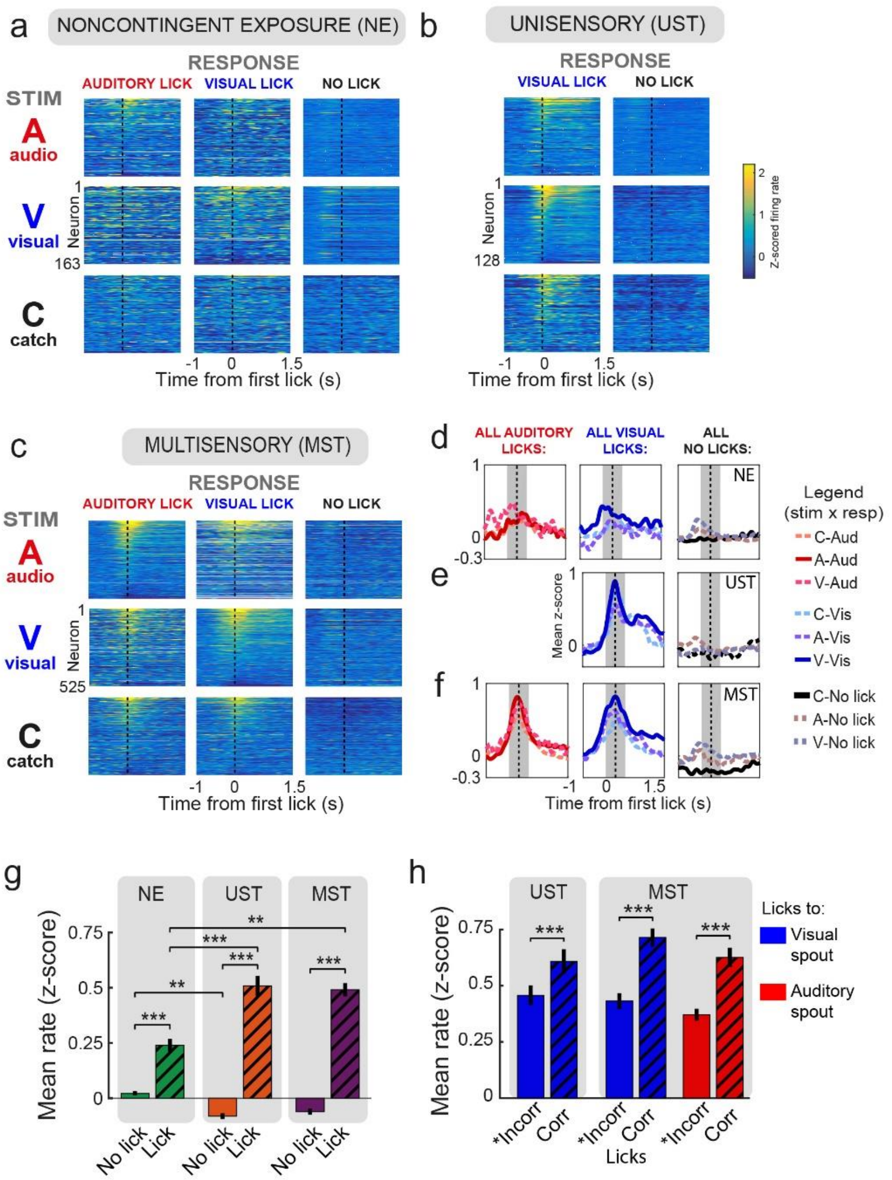


**Extended Data Figure 3: Licking activity modulates V1 activity across training conditions (Task A).** We computed the average z-scored activity across all recorded neurons in V1 aligned to the first lick for 9 stimulus-response combinations: three stimulus type conditions (A=auditory, V=visual, and C=catch – i.e. no change) and three response options (lick to visual spout, lick to auditory spout, and no lick). a-c) In NE mice, all conditions with licking (left 6 heatmaps) showed slight modulations of activity around licks, which were absent in the three conditions without licking (right 3 heatmaps). This lick-aligned modulation, however, was much more prominent in MST and UST mice (panels **b** and **c**, respectively). For trials in which there was no lick, activity was aligned to the median response latency from all other trials. Conditions for which not enough trials were present to compute a reliable mean z-score for that neuron (fewer than 3 trials), were omitted from the heatmap. Therefore, trial-averaged estimates for licks to the auditory spout are absent in UST animals (trained on vision only), as animals rarely responded to the never-rewarded auditory lick spout. d-f) All conditions with licking (left 2 panels for each row) show activity modulations hundreds of milliseconds before and after lick-onset. Each plot combines the heatmaps of three conditions (taken per column) and shows the Z-scored firing rate averaged over neurons. I.e. the left panel shows the average for the three columns (**d**: NE; **e**: UST; **f**: MST). Lines show mean across neurons. As in *b*, not enough events were available to calculate licks to the auditory lick spout in UST animals. g) Licks evoked consistently higher V1 firing activity than no-licks for all cohorts ($p < 0.001$) in the time window around lick onset (-300 ms to +300 ms relative to lick onset, gray-shaded patch in d-f). Lick modulation was stronger for trained cohorts versus naive mice (UST vs NE, MST vs NE, *$p<0.05$, **$p<0.01$, ***$p<0.001$). h) Correct hits were associated with higher V1 firing activity than incorrect licks in all trained cohorts ($p < 0.001$). *Incorrect licks include both false alarms and mistaken licks to opposite spout (e.g. visual stimulus, lick to auditory lick spout). Conditions are separated to allow comparing between licks to the same spout.

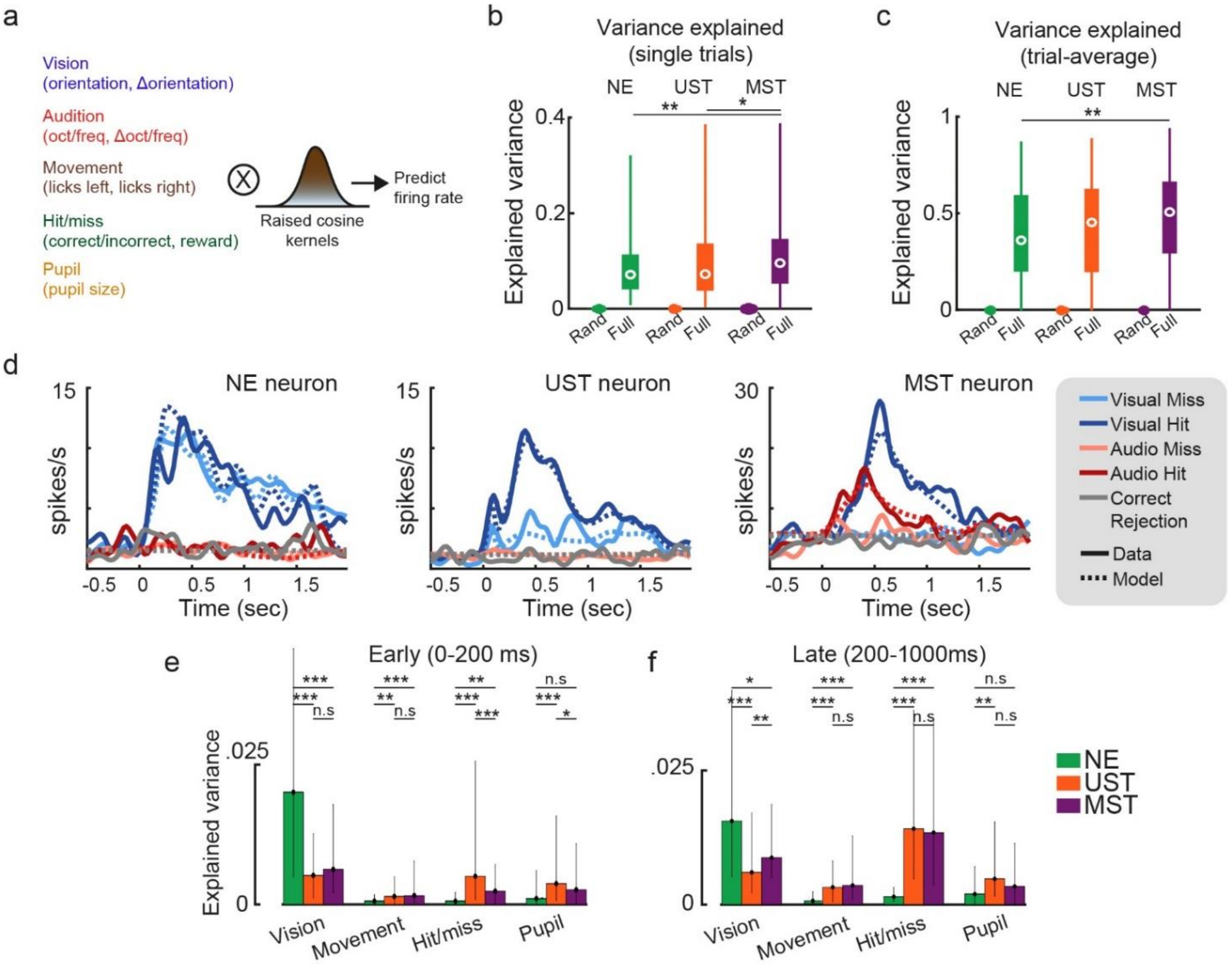


**Extended Data Figure 4: Application of the generalized linear encoding model to explain variance in firing rates from different cohorts (Task A).** a) To dissociate the contribution of sensory, task, movement, and arousal variables we built a kernel-based regression model[22,37,38] where stimulus variables (visual and auditory features and amount of change), as well as report variables (hit/miss), motor variables (licks left and right) and pupil size, were included as predictors of firing rate. Binary task variables (all but pupil size) were convolved with raised cosine basis functions that spanned the relevant time window to model transient firing rate dynamics. b) We constructed two models. The first model operated as a null model and consisted of a random predictor only (Rand). The second model included all predictors as in **a** (Full). We quantified the model performance in two ways. First, we computed the cross-validated explained variance (EV) over all single-trial firing rates for all neurons (n=116 NE neurons, n=128 UST neurons, n=272 MST neurons). The full model always explained significantly more variability than the random model (all p<1e-20), and explained variance was slightly higher for V1 neurons from MST mice than NE or UST neurons (NE vs UST, p>0.05; NE vs MST, p=0.007; UST vs MST, p=0.033). White dots show the median and the width of the violin plot shows neuron density. c) We also quantified model performance by computing the EV of the firing rate averaged over the five stimulus x choice conditions that captured nearly all trials[20,38]. Again, the full model explained significantly more variability than the random model (all p<1e-33), while variability across task versions was comparable but higher for MST than NE neurons (p=0.001). d) Model fits for three example neurons (one V1 neuron from each task version) show that predicted and actual firing rates closely overlap and that both sensory-driven activity (example #1), as well as report-related activity for both visual and auditory hits (examples #2-3), are captured by the model. The five stimulus-response combinations that had the most counts are plotted (trials with false alarms and licking to the incorrect spout are omitted). e) We computed the variance of the firing rate as explained by each subset of predictors for each of the task versions over time (cf. Figure 2g). During the early post-stimulus window (averaging EV over 0-200 ms) visual predictors explained more variance in NE than UST and MST mice. Bar and errors show median and interquartile range. *p<0.05, **p<0.01, ***p<0.001. f) Same as **e**, but for the late window (200-1000 ms). The hit/miss predictor now explains more variance than during the early period and explains more variance in both UST and MST than in NE mice (p<0.001). Therefore, knowing when licks occurred and whether the trial was a hit or not, contributed to predicting late V1 firing in UST and MST mice. Moreover, visual predictors continue to make strong contributions in the late phase across the three training cohorts.

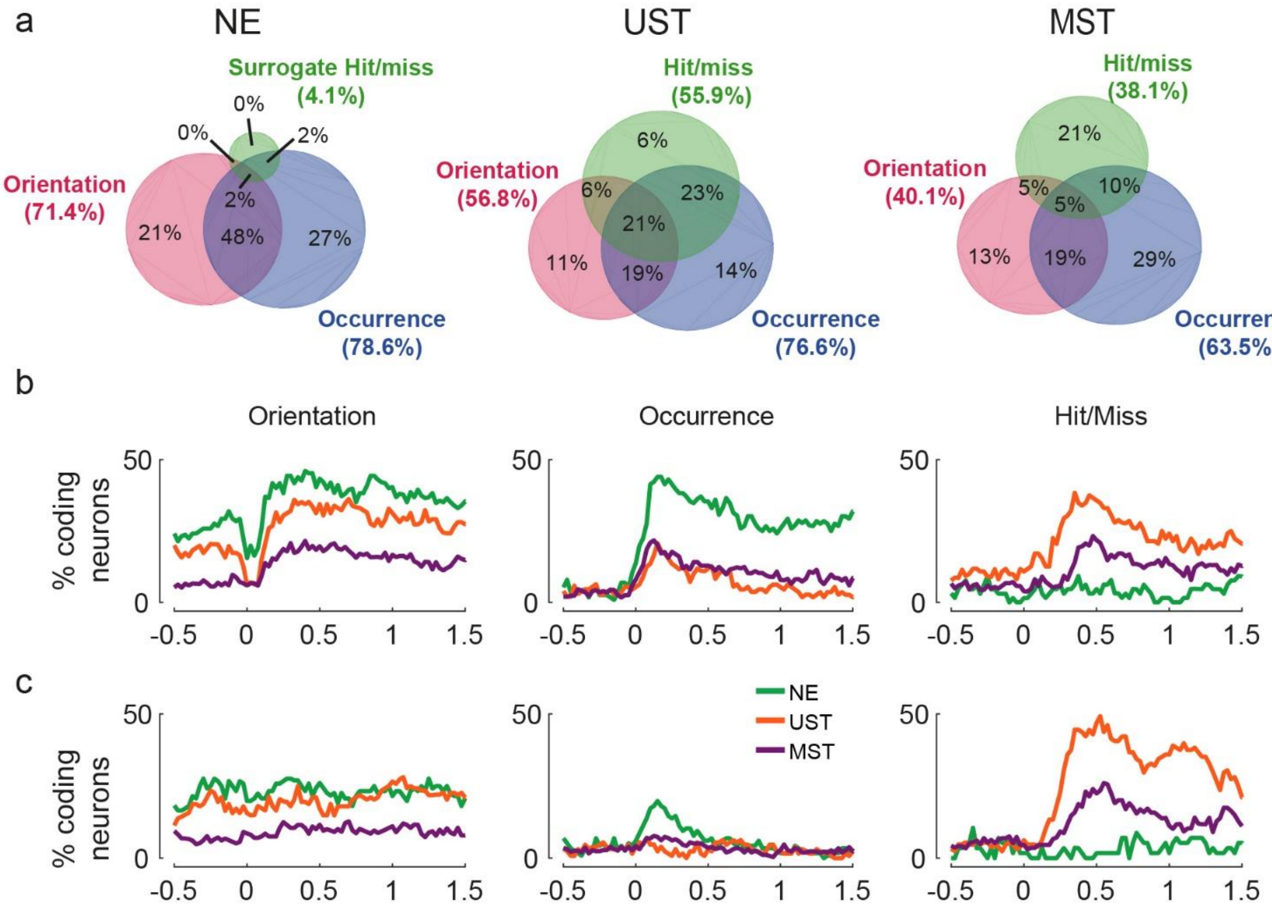


**Extended Data Figure 5: Report-related coding in firing patterns recorded from deep layers of trained animals (Task A).** a) The Venn diagrams show for each training cohort the percentage of neurons encoding orientation (grating after stimulus change), occurrence (presence of a visual change or not), or hit/miss as established with ROC analysis. Neurons were classified as coding if their AUC score was significant for at least three time bins (=75 ms, permutation test) during the 0 to 1000 ms window after a stimulus change. Only maximum visual change trials were used. Similar to our encoding model (Ext. Data Fig. 4) and in line with the averaged z-scored activity (Fig. 2d-f), report-related coding was only substantially present in UST and MST mice, while the ratios of orientation and visual occurrence coding neurons were comparable across cohorts. Shown are percentages out of all coding neurons; percentage of non-coding neurons per cohort: NE: 15.5%; UST: 13.3%, MST: 35.6%. Each value inside the Venn diagrams indicate the percentage of neurons falling in the indicated portion of the diagram, and summing to 100%. Conversely, the sum percentages indicated outside the Venn diagrams sums to more than 100% as neurons can show different forms of encoding. b) Percentage of significant neurons encoding selected variables over time for visual changes of maximal saliency. The strength of encoding (AUC values above shuffled) gave similar results as the fraction of coding neurons (shown here). c) Same as *b*, but for threshold visual changes.

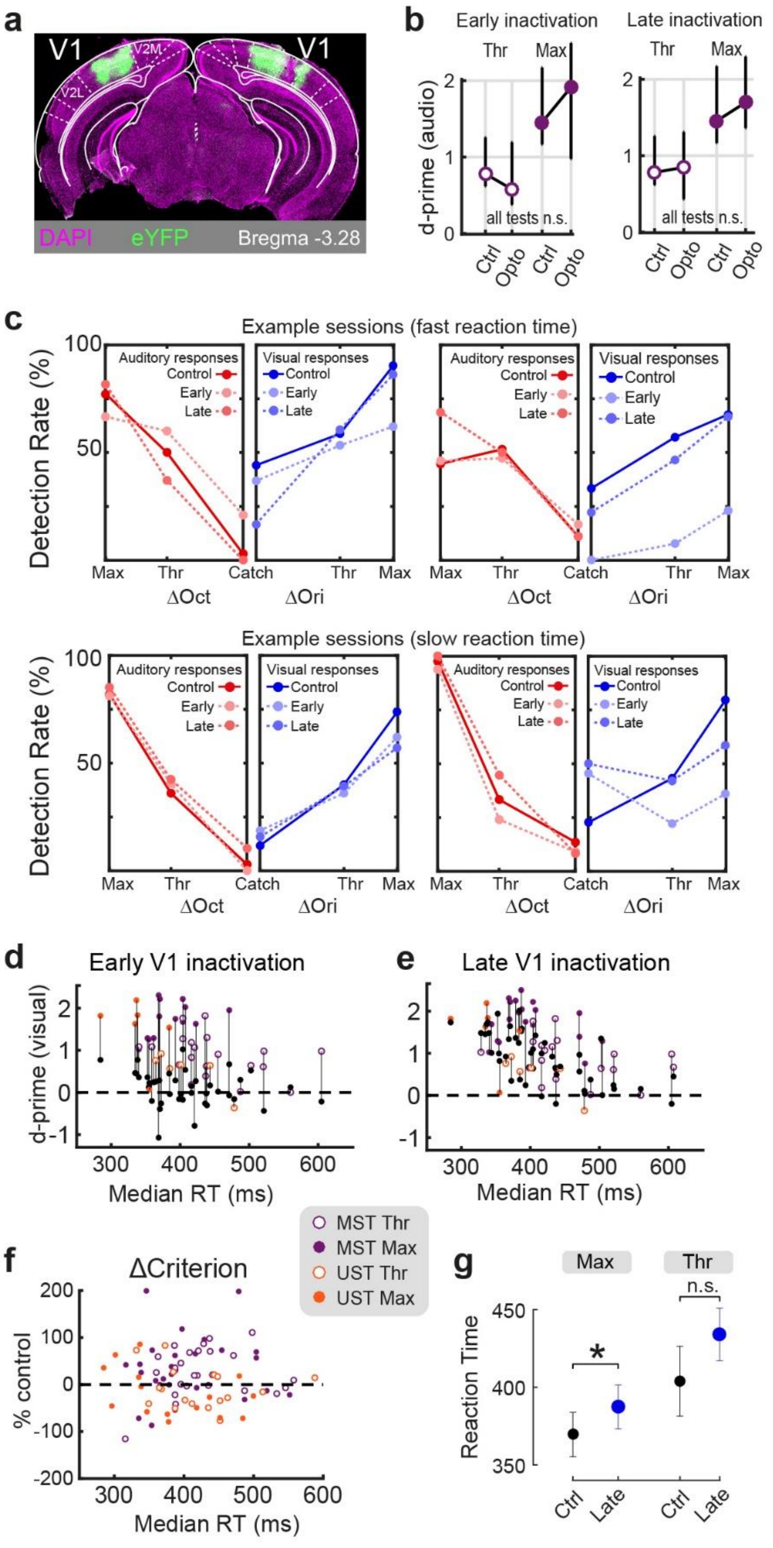


**Extended Data Figure 6: Detailed characterization of early and late V1 inactivation in task A.**

a) Coronal histological section revealing localized bilateral expression in V1. V2L: lateral secondary visual cortex. V2M: medial secondary visual cortex.

b) Neither early nor late silencing affected auditory discrimination performance (d-prime) for both threshold and maximum saliencies and across UST and MST cohorts (all p>0.1).

c) Behavioral detection rates for two example sessions with fast reaction times (top; median visual hit reaction time 359 and 362 ms) and two sessions with slow reaction time (bottom; 435 and 463 ms). Late silencing affects slow sessions, but not fast sessions. Example sessions are all from MST mice.

d) Scatter plot of visual d-prime on control (colored) and photostimulation trials (black). One data point is one session. Data points from the same session are connected with a line to visualize the reduction in d-prime.

e) Same as **d** but for late silencing. Note how sessions with short reaction time are proportionally less affected than sessions with long reaction time. Quantification of this effect as the percentage reduction in d-prime is in Fig. 3i,j.

f) We tested whether late inactivation could affect motivation by changing the criterion parameter in our signal detection framework (see Methods). The reduction in visual criterion by late photostimulation was not significantly correlated to the median reaction time on control trials in the same recording session (p=0.291).

g) As late photostimulation partially reduced hit rate for visual changes in MST mice depending on reaction time (Fig. 3h,j), some visual changes were still detected. Comparing the reaction time of those visual hits with control visual hits revealed that late photostimulation delayed reaction times for maximal visual changes in MST mice (MST – *max*: p=0.024; MST – *thr*: p=0.177). Traces show mean ± SEM across visual hits. *p < 0.05.

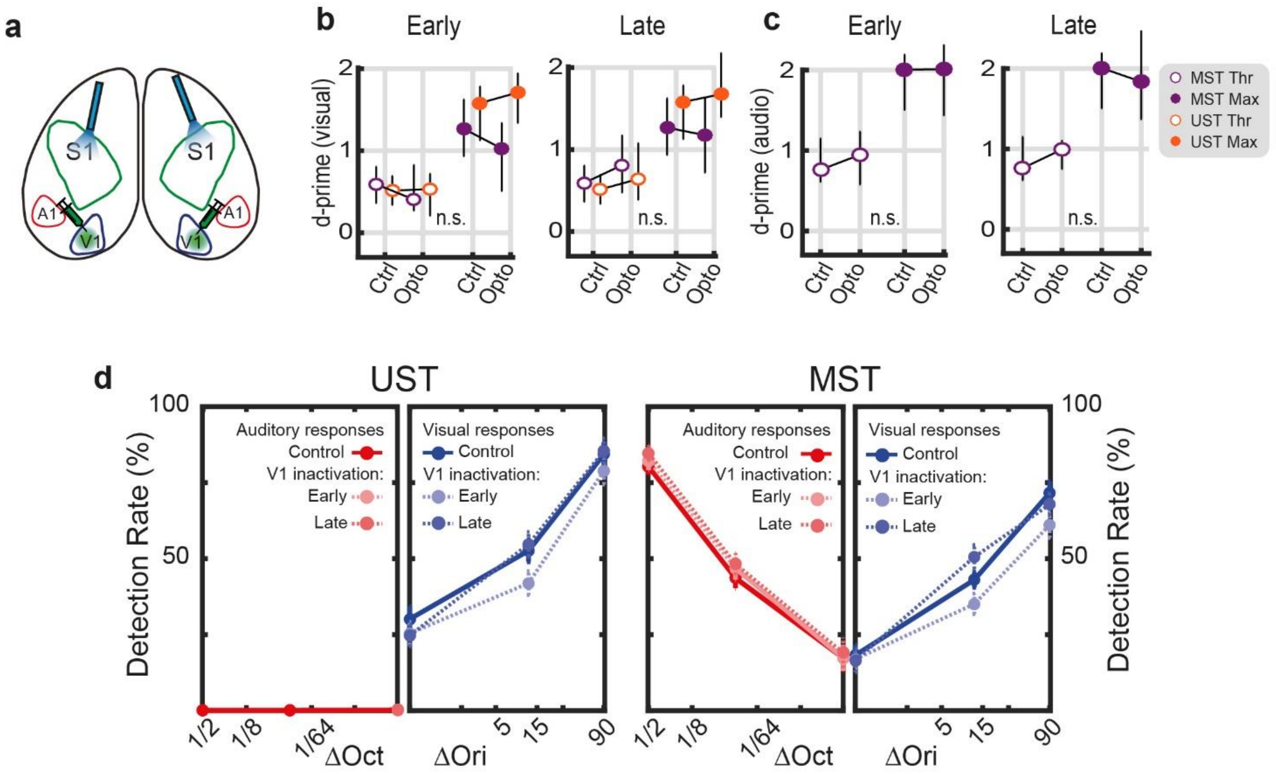


**Extended Data Figure 7: Illumination of control area S1 in task A has no behavioral effects.** a) Control experiment with positioning of the optical fiber over uninfected S1. b) D-prime across visual conditions. Neither early nor late S1-illumination significantly affected visual detection performance (all p>0.05). c) Same as **b**, but for auditory conditions. Neither early nor late S1-illumination significantly affected auditory detection performance (all p>0.05). d) Behavioral response rates for UST (left) and MST (right) mice for control, early and late S1-illumination trials.

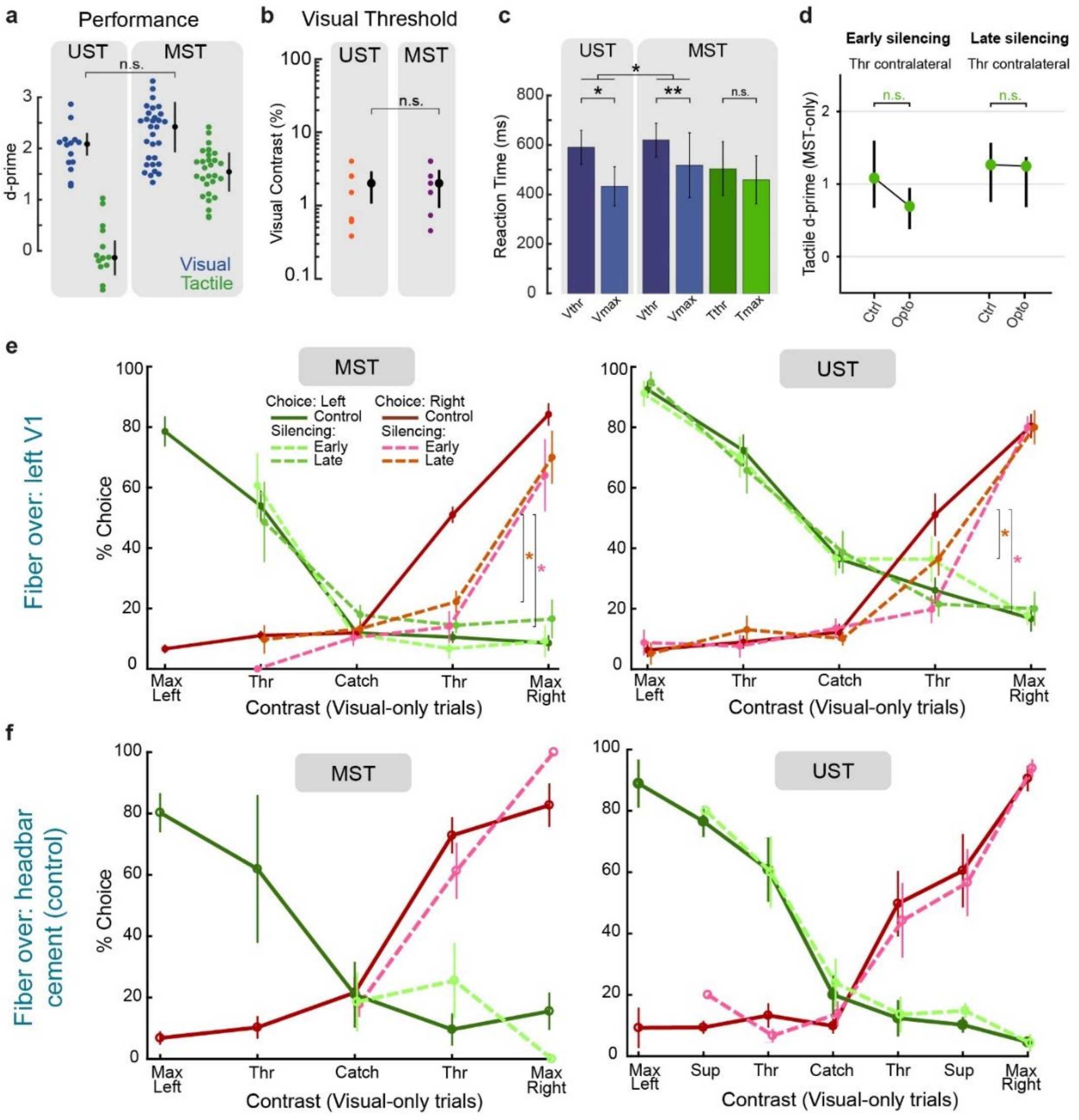


**Extended Data Figure 8: Effects of optogenetic V1 silencing on visuotactile behavior (task B). a)** D-prime at maximum saliency for visual and tactile detection. Each dot represents one session and either right or left side detection performance. Visual performance was comparable for UST and MST mice (p=0.147). Note that null d-prime for tactile detection is expected for UST mice. **b)** Visual contrast detection thresholds were comparable for UST and MST mice (p=0.701). Computed for each mouse from psychometric fit, for both right and left side detections. **c)** Median reaction times for each rewarded condition for threshold and maximum levels of saliency (right and left sides pooled together). Visual reaction times were significantly shorter for UST compared to MST (p=0.011, both saliencies pooled. *: p<0.05; **: p<0.01). Note that tactile and visual reaction times were similar, ruling out the possibility of a sequential detection strategy where one modality would be sampled before the other one. **d)** D-prime of contralateral tactile detection at threshold saliency. V1 silencing did not affect tactile performance (Early silencing: p=0.264; Late silencing: p=0.667). Thus, for MST, silencing late V1 activity impaired visual but not tactile detection, indicating that late activity per se is not required for licking behavior. **e)** Psychometric curves for experiments with left hemisphere V1 silencing (same experiments as Fig. 4d), for visual-only trials. Points not shown were not part of the experimental protocol. Since monocular stimuli were used and left hemisphere V1 was silenced, potential effects were expected for stimuli on the right side (contralateral) but not on the left side (ipsilateral). (Compared to control: MST Early: p=0.0469; MST Late: p=0.0469; UST Early: p=0.0313; UST Late: p=0.0156; *: p<0.05). Note that late silencing for UST mice had a significant effect on the percentage of right choices, but no effect on the corresponding d-prime. **f)** Psychometric curves for control experiments where the optic fiber was placed above the mouse headbar cement and therefore not above V1 (all not significant).

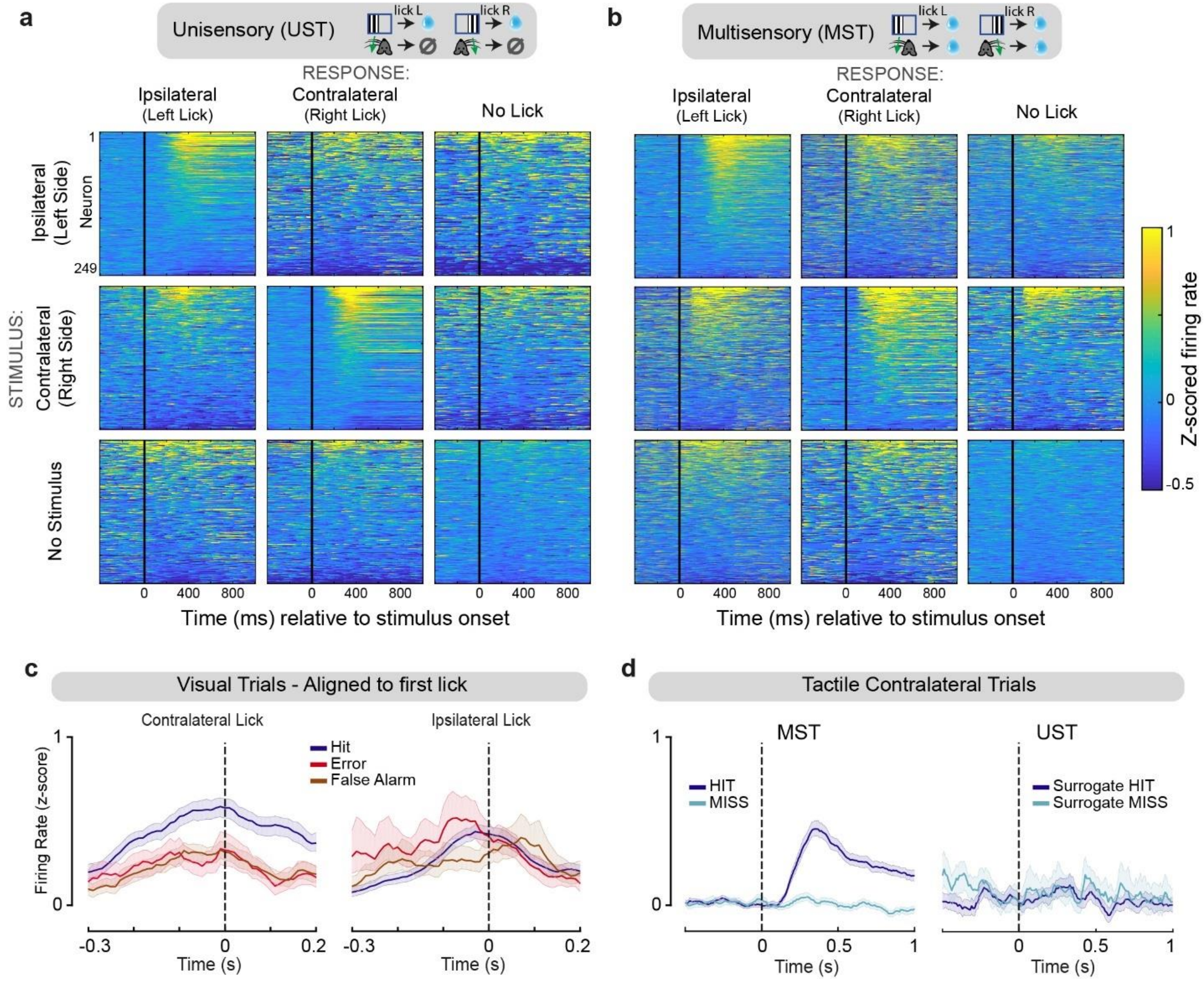


**Extended Data Figure 9: V1 neuronal responses across stimulus-response conditions in the visuotactile detection task (Task B) encode a combination of sensory and non-sensory factors during the late time window. a,b)** Average z-scored activity for all recorded left-hemisphere V1 neurons for each of all 9 possible visual-only stimulus-response combinations for UST **(a)** and MST **(b)** trained mice. Neurons for which no more than three trials were present in the given condition were omitted. For each condition, neurons were sorted according to their mean z-score between 50 and 500 ms. **c)** Average z-scored firing rate of responsive V1 neurons during visual-only trials, for three different conditions eliciting the same licking response (left: contralateral lick, right: ipsilateral lick), showing that late V1 activity cannot be explained by licking alone (see also Ext. Data Fig. 3). Activity was aligned to the first lick in the response window. Licks made to the wrong side were termed "errors". Same neurons as in Fig. 4c. Shaded area: bootstrapped 95% confidence intervals. **d)** Average z-scored firing rate of responsive V1 neurons during contralateral tactile trials, split by choice, for UST and MST mice. For UST, surrogate hits correspond to licks to the same side of the tactile stimulus (although unrewarded) and surrogate misses correspond to trials without licks. Late activity was present only in MST Hits, indicating that the same stimulus and the same behavioral response triggered late activity in a context-dependent manner. Shaded area: bootstrapped 95% confidence intervals.

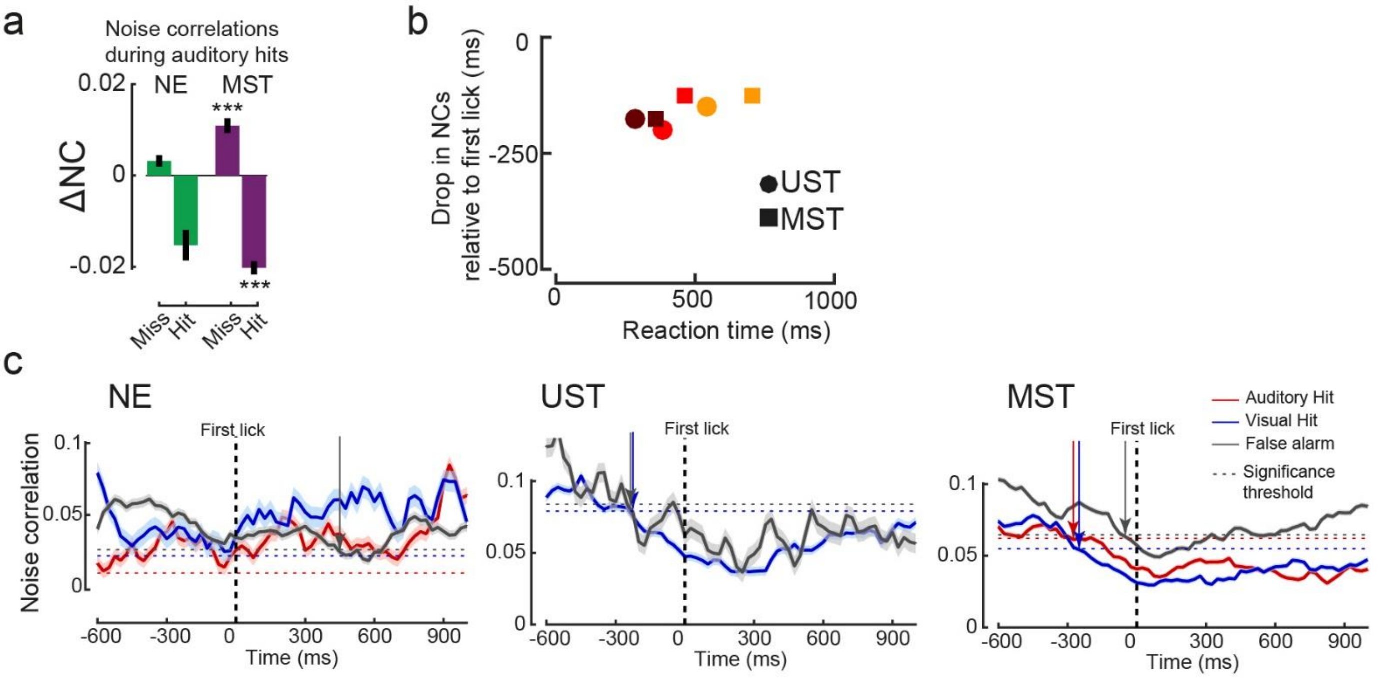


**Extended Data Figure 10: Noise correlations in auditory trials and latency of the drop in noise correlations during visual trials in task A. a)** Decrease in noise correlation (NC) of V1 cell pairs with respect to baseline for auditory hits and misses across cohorts (same conventions as Figure 5c). For UST mice there were too few auditory hits to compute noise correlations. In MST mice, noise correlations decreased during hits (p=7.3e-16) and increased during misses (p=1.8e-9). The fact that NCs also decrease in auditory hits indicates that the drop in NCs in the visual cortex is not specific to visual trials, and possibly is a more general mechanism that may subserve decision making. **b)** Drop in NCs during visual trials, but relative to the first lick for each tertile of reaction times for UST and MST. Same as Fig. 5f, but aligned to reaction time. No significant correlation is found (p=0.09), in contrast with Fig. 5f, indicating that the drop in NCs precedes reaction time by a relatively constant time lag. **c)** Noise correlations (NC) of V1 cell pairs over time aligned to the first lick for the different trial types and cohorts. Dotted line shows the threshold for a significant drop in NCs with respect to baseline (-1000 to -500 ms relative to first lick). Noise correlations decreased most for visual hits in visually trained mice (UST and MST), but not in NE mice. For UST mice there were too few auditory hits to compute noise correlations.
